# The structural scaffold of the TPLATE complex deforms the membrane during plant endocytosis

**DOI:** 10.1101/2024.10.07.616965

**Authors:** Julia M. Kraus, Michaela Neubergerová, Alvaro Furones Cuadrado, Neeltje Schilling, Dominique Eeckhout, Nancy De Winne, Eveline Van De Slijke, Michael Vandorpe, Klaas Yperman, Evelien Mylle, Marcus Fislage, Geert De Jaeger, Roman Pleskot, Daniel Van Damme

**Author notes:** Corresponding and senior authors. equal contribution (Julia M. Kraus, Michaela Neubergerová).

## Abstract

Eukaryotic cells maintain homeostasis of their outer membrane by controlled internalization of lipid and protein constituents via endocytosis^1^. Endocytosis is evolutionary conserved and utilizes similar structural folds. How these structural folds are combined into proteins and protein complexes however differs between eukaryotic kingdoms^2^. The TPLATE complex in plants is an evolutionary ancient protein module that combines several endocytic folds into a single octameric protein complex^3–5^. Its molecular architecture, lipid-nucleated condensate formation, and its requirement for clathrin cage curvature revealed its function in endocytosis initiation in plants^6–8^. Mechanistic understanding of how this complex drives membrane deformation during plant endocytosis is, however, lacking. Here, we used an integrative structural approach to obtain a precise molecular structure of the TPLATE complex. In addition, our approach allowed visualizing the structural flexibility that hallmarks this enigmatic complex. We prove that the intrinsic structural flexibility is required for its functionality and membrane recruitment. The membrane binding interface consists of several domains with differential lipid preferences. Finally, we show that the crescent shape of the structured part of the complex is sufficient for membrane curvature generation. Our mechanistic insight answers the long-standing question of how plants execute endocytosis without cytoskeletal-based force generation.

## Main text

Endocytosis is an essential multi-step process in all eukaryotic cells to internalize lipids, transmembrane proteins and other cargo from the plasma membrane (PM)^1^. While its execution shares common structural domains and even conserved adaptor protein complexes, different kingdoms mechanistically fine-tuned the process to their specific needs^2^. Endocytosis predominantly uses the adaptor protein complex 2 (AP-2) in combination with the scaffolding molecule clathrin^9^. Clathrin-mediated endocytosis (CME) involves condensate formation, which creates micro-environments that allow successful cargo recognition^7,10–12^ alongside clathrin recruitment^7,13,14^. Next to AP-2, plants require an additional adaptor protein complex called TPLATE complex (TPC) to execute endocytosis^3^. TPC is evolutionary ancient, belongs to the heterotetrameric adapter complex containing a coat (HTAC-CC) family, and shares its origin with AP-2 and coat protein complex I (COPI)^15^. Besides plants, TPC has so far only been experimentally characterized in the slime mold Dictyostelium^4^. TPC has been lost in animals and yeasts, which explains its late discovery^4,5,15^. TPC is an octameric complex with a molecular weight of roughly one megadalton. Several of its subunits contain a considerable percentage of intrinsic disorder^2^. It is essential for plant life as single subunit knockouts cause pollen lethality^3^. Two of its subunits, namely AtEH1/Pan1 and AtEH2/Pan1, which both have a disordered content of more than 60 percent, form condensates and recruit both early and late endocytic players, including clathrin^7^. TPC is recruited to the PM as an octameric complex^16^ and features multiple membrane binding domains, as well as cargo recognition domains^6,17,18^. Initial structural insight into the arrangement of all eight subunits was achieved by integrating protein-protein interaction assays, chemical cross-linking mass spectrometry (XL-MS), and computational modeling^6^. Here, by combining electron microscopy (EM), XL-MS and computational modeling, we provide a comprehensive integrative structure of TPC, which is at par with other HTAC-CC structural models such as the COPI complex or the clathrin-AP-2 complex^19–21^. Our structural model visualizes the functional interplay between structured and disordered parts of TPC, describes the PM-binding interface of the complex and its domains in detail, and elucidates how the structured parts provide the force required to bend the PM.

### Determining a TPC structure

To obtain the complex, TPC was purified from Arabidopsis PSB-D cell suspension cultures using tandem-affinity purification (Extended Data Figure 1 a & b). One TPC assembly features a net charge of −133 at physiological pH, which results from 1000 negatively and 867 positively charged residues. Together with the disordered nature of several subunits of the complex, this caused significant air - water interface problems. We could not determine a condition where these problems were mitigated and therefore this did not allow cryo EM-based visualization of the complex. However, negative stain (ns-)EM analysis presented a good particle count and distribution (Figure 1 a). The complex displayed preferred orientations on the EM grid, resulting in a limited number of different 2D classes (Figure 1 b, Extended Data Figure 1 d & i).

**Figure 1.**
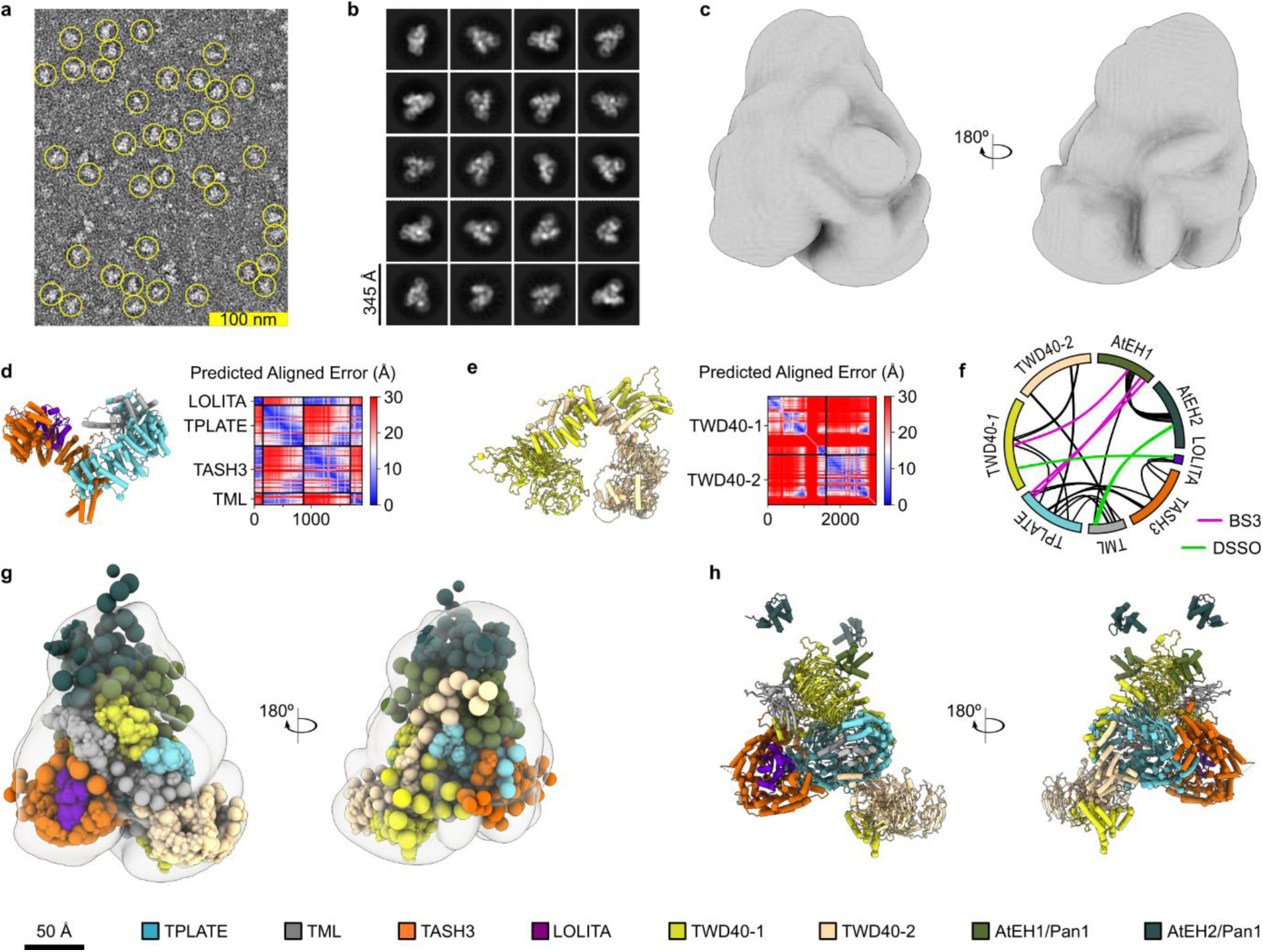
The structure of the TPLATE complex is determined via integrative modeling. a) A representative negative stain (ns)EM micrograph of purified octameric TPC from Arabidopsis cell culture cells. Good quality particles are circled in yellow. The scale bar equals 100 nm. b) A selection of 2D classes depicting the different orientations of TPC that were observed. The scale bar equals 345 Å. c) Two views of the 3D map of TPC, 180 degrees rotated and reconstituted from 156.997 selected particles to a resolution of 17 Å and then gaussed to approximately 20 Å to avoid over estimation. d) AlphaFold2 prediction of the TPC inner core consisting of the TASH3 trunk domain (orange), LOLITA (purple), the TPLATE trunk domain (teal) and the TML longin domain (gray). The predicted aligned error plots show that AlphaFold2 is confident about the interaction domain interfaces. e) TWD40-1 (yellow) and TWD40-2 (beige) sub-complex prediction with AlphaFold2. The predicted aligned error plot indicates a confident prediction of the subunits, as well as of the arch closure of the two subunits. The relative position of both propeller domains is however uncertain. f) Visualization of MS identified inter-molecular cross-links between the TPC subunits mapped on a circular representation of the eight subunits. BS3 and DSSO share structural information provided by the 38 black cross-links. BS3 (pink) and DSSO (green) also provide unique structural information. The dataset consists of 45 high confidence inter-molecular cross-links in total. g) Multi-scale TPC model obtained with the data provided by panels C to- F via integrative modeling. The model represents a centroid structure, i.e. the most similar structure to all other structures in the top-scoring cluster. nsEM is shown in transparent gray. h) The centroid model shown as a ribbon diagram for the well-structured TPC domains described as rigid bodies in integrative modeling.

We reconstructed 3D maps from three independent purifications in two different buffer systems. All independently reconstituted maps yielded the same dimensions, the same shape, the same resolution, the same preferred orientation and a correlation of more than 82 % (Extended Data Figure 1 c). The final map achieved an approximate resolution of 17Å (Extended Data Figure 1 g). We subsequently took advantage of an integrative approach to achieve a high-resolution structure, taking into account that such an approach enables generating a more complete, accurate and precise model compared to those that are based on a single method^22^. To generate a structural model of TPC, we combined the map acquired by nsEM with chemical cross-link data as position restraints for the integrative modeling^23^ (Figure 1 c & f). The chemical cross-links that we used included data using an MS-cleavable cross-linker, DSSO, as well as previously published BS3 cross-linking data (Figure 1 f)^6^. The majority of the DSSO cross-links were consistent with the BS3 dataset. However, some unique DSSO cross-links provided new connections between TPC subunits, particularly between LOLITA and the N-terminal part of TWD40-1 and between the TML μ-homology domain (μHD) and the C-terminal part of AtEH2/Pan1 (Figure 1 f). Highly reliable structural models of individual subunits and partial sub-complexes generated by AlphaFold^24–26^ were used as input for the well-structured parts of the complex (Figure 1 d & e, Extended Data Figure 2 a).

Integrative modeling resulted in a structural model of TPC with a precision ranging from 5 to 50 Å and with an average precision of 31.5 Å (Extended Data Figure 2 & Extended Data Figure 3). Including all parts of the complex in the structure-building process allowed us to analyze the precision of different TPC parts at the given resolution and positional restraints, and to identify the regions of low precision, which are likely representing flexible parts of the complex. The highly flexible parts of the complex are represented mainly by the highly disordered AtEH1/Pan1 and AtEH2/Pan1 subunits. In addition, we observed high flexibility for the TML linker region, the μHD, the SH3 domain of TASH3 and the C-terminal all-alpha domain of TWD40-1 (Extended Data Figure 3). In contrast, the structural model of the hexameric core of the TPC, consisting of LOLITA, TPLATE, the TASH3 trunk domain, the TML longin domain, the β-propeller domains together with the solenoid domain of the TWD40-1 and TWD40-2 subunits, displayed a high to medium precision, suggesting low flexibility within the complex. Consistently with our previous structural model of TPC^6^, the complex has an overall ellipsoidal shape with the TPLATE subunit forming an interaction hub connecting many other TPC subunits (Figure 2 g & h).

**Figure 2.**
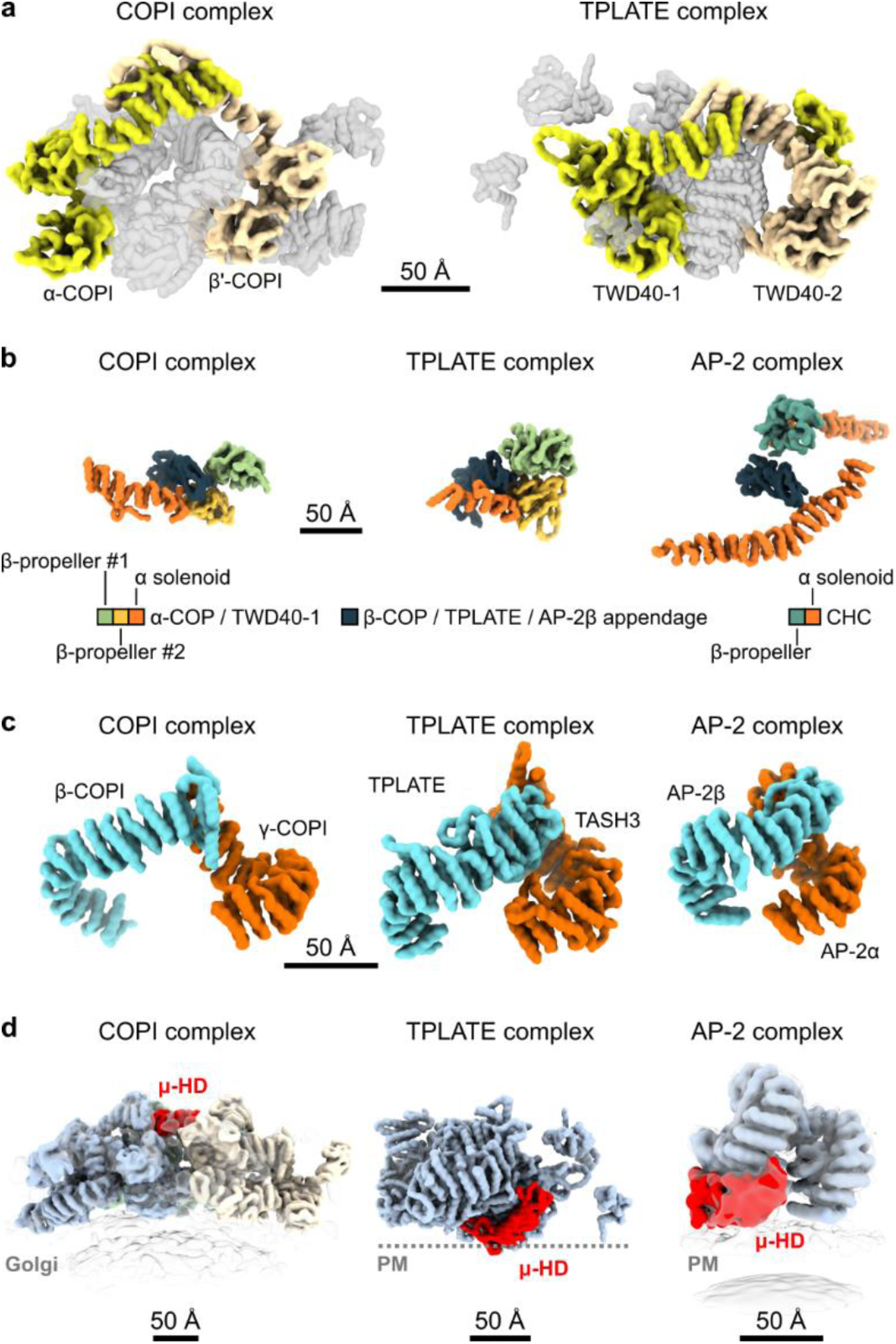
TPC positions structural domains intermediate to both AP-2 and COPI. The backbones of the structures without the side chains are visualized as a surface in UCSF ChimeraX. a) Structural comparison of TPC and COPI (PDB code 5NZR). Visually aligned complexes share two subunits arching the core complex. These subunits have been lost in AP-2. TWD40-1 relates to α-COP (both yellow), while TWD40-2 relates to β’-COP (both beige). The subunits are structurally conserved in their domain organization, with two N-terminal β-propellers followed by an α-solenoid that closes the arch. b) The appendage domains of TPLATE, β-COP and AP-2β (dark blue) all interact with α-solenoids (orange) and β-propeller domains. In TPC and COPI, the two β-propeller domains (light green and yellow) originate from the arching subunits TWD40-1 and α-COP. This interaction is structurally conserved. In AP-2 the mode of interaction structurally differs and the β-propeller domain (turquoise) belongs to clathrin heavy chain. For COPI the leaf structure (PDB code 5NZR) was used. For AP-2 the clathrin with AP-2β appendage structure (PDB code 6YAI) is visualized. c) The large core subunits, TPLATE, β-COP and AP-2β (all turquoise) and γ-COP, TASH3 and AP-2α (all orange) of each complex (COPI leaf PDB 5NZR, AP-2 open conformation PDB 6YAH) all connect at their C-termini, while their N-termini are separated from each other. The distance between the N-termini is largest for COPI and smallest for AP-2, while TPC positions itself between the two complexes. d) Visualization of the position of the μHD of each complex (red). In COPI, it is distal of the membrane and aids polymerization in some COPI linkages, like in the one displayed (PDB code 5NZU). In both AP-2 (PDB code 6YAH) and TPC the μHD establishes direct PM contact. For TPC the PM is presented as a dotted line. Scale bars equal 50 Å.

To generate a model of the TPC molecular structure that was not dependent on the input data, we utilized the AlphaFold3 (AF3) server^26^. The token limit only allowed modeling a truncated hexamer containing the TASH3 trunk domain, LOLITA, the TPLATE trunk domain, the TML longin domain, and the β-propeller domains, along with the solenoid domain of the TWD40-1 and TWD40-2 subunits (Extended Data Figure 4 a & b). The resulting AF3 model of the truncated TPC hexamer closely aligns with the integrative model. However, there are clear differences in the position of the TASH3-LOLITA pair between the two models and the position of the TASH3-LOLITA pair in the AF3-generated model violated several cross-link restraints (Extended Data Figure 4 c). These results are in line with the limitation of artificial intelligence-based protein structure prediction^27^ and highlight the strength of the integrative modeling approach^22^. However, it is possible that the AF3 model represents a different conformation of the complex.

The structural model that we obtained with the integrative modeling approach subsequently allowed us to compare TPC to other members of the HTAC-CC family.

### TPC is intermediate to HTAC-CC members

The heteroheptameric COPI and the heterotetrameric AP-2 complex have both been structurally resolved to the level of their vesicle coats^19,20^. ConSurf^28^ analysis (Extended Data Figure 5) revealed strong conservation of these two complexes throughout evolution, allowing us to compare the available non-plant structural models to the TPC structure (Figure 2). The TPC structure corroborates the previous phylogenetic analysis, indicating that COPI and TPC originated from the root of the tree of the HTAC-CC family^4,15,29^. While both TPC and COPI have two subunits arching their core, AP-2 has lost these subunits^4^. These arching subunits between TPC and COPI are conserved down to their domain organization. They both have two N-terminal β-propeller domains followed by a large α-solenoid where they interact with each other to close the arch (Figure 2 a). Consistently with the shared evolutionary origin of the HTAC-CC family, the appendage domains of TPLATE, β-COP and AP-2β all interact with β-propeller domains. In case of TPC and COPI, the interacting β-propeller domains are provided by the arching subunits TWD40-1 and α-COP and this interaction is structurally nearly identical (Figure 2 b). The AP-2β appendage interacts with a β-propeller provided by clathrin heavy chain (CHC), however the mode of action of this interaction is different compared to the other complexes (Figure 2 b). These findings support the closer relation of TPC to COPI than to AP-2. When comparing the core subunits of each complex (Figure 2 c), the two large TPC subunits are positioned in an open conformation. However, this open conformation of TPC lies in between the open conformations of COPI and AP-2. COPI requires its μHD for complex polymerization in linkages II and III at the distal end of the vesicle^19,30^, while in both AP-2 and TPC this domain is essential for lipid interactions^6,31^. Consistently, the μHD of AP-2 is located close to the vesicle^20^ and previous data also positions the μHD of TPC at the membrane interface^6^. Altogether, our structural findings are, therefore, consistent with the phylogenetic analysis of the HTAC-CC family, while the role of the μHD in the different adaptor protein complexes presents an intriguing functional diversification. Since COPI and AP-2 both feature a clear membrane interacting interface, we next analyzed the TPC-membrane interaction.

### Specific domains bind the membrane

To analyze the membrane binding capacity of purified TPC *in vitro*, we applied the complex to large unilamellar vesicles (LUVs) with various lipid compositions and performed flotation assays. TPC differentially bound to vesicles with negatively charged lipids, and the strongest interaction, resulting in the lowest amount of TPC in the pellet fraction, was observed for phosphatidic acid (PA) containing vesicles. Uncharged lipids did not result in a clear interaction (Figure 3 a). These results were confirmed via LUV tubulation experiments. (Figure 3 b). LUVs without TPC did not tubulate under our experimental conditions (Extended Data Figure 6) and in the presence of TPC LUVs containing uncharged lipids presented nearly no tubulated vesicles. Tubulation of charged LUVs, however, correlated with TPC concentration (Figure 3 b). Coarse-grained molecular dynamics (MD) simulations with a proxy lipid bilayer to the plant PM confirmed our *in vitro* experiments by showing that TPC clusters anionic phospholipids at the membrane binding interface (Figure 3 c, Extended Data Figure 7). MD simulations were performed with the hexameric model as the highly disordered AtEH/Pan1 subunits hampered with modeling convergence (Figure 3 c). The hexameric model resided stably on the membrane in 4 out of 5 simulation repeats (Extended Data Figure 7 a) with the same interaction mode (Extended Data Figure 7 b) corresponding to the positively charged patches on the TPC surface (Figure 3 c). The phosphatidylinositol 4-phosphate (PI4P) preference of the μHD of TML was described previously by MD simulations using the separated domain, as well as by *in planta* experiments^6^. Both EH1.1 and EH1.2 domains of the AtEH1/Pan1 subunit have a strong binding capacity for PA^17^. By including other subunits in MD simulations as part of the complex, we observed that, consistently with the single domain MD simulations, the TML μHD preferred PI4P. Additionally, we found that the N-terminal part of TASH3 also displayed a strong preference for PI4P. In contrast, LOLITA, TWD40-1 and TWD40-2 preferred phosphatidylinositol 4,5-bisphosphate (PI4,5P2) (Figure 3 c). The MD simulations using the hexameric complex did not identify a domain with strong interaction with phosphatidylserine (Extended Data Figure 7 c & d), nor a domain with a preference for PA, indicating that the AtEH/Pan1 subunits are responsible for the observed binding to PA in our flotation assays. Next, we focused on the β-propeller domains of the TWD40-1 subunit since they indicated an uncharacterized PI4,5P_2_ preference of the TPC.

**Figure 3.**
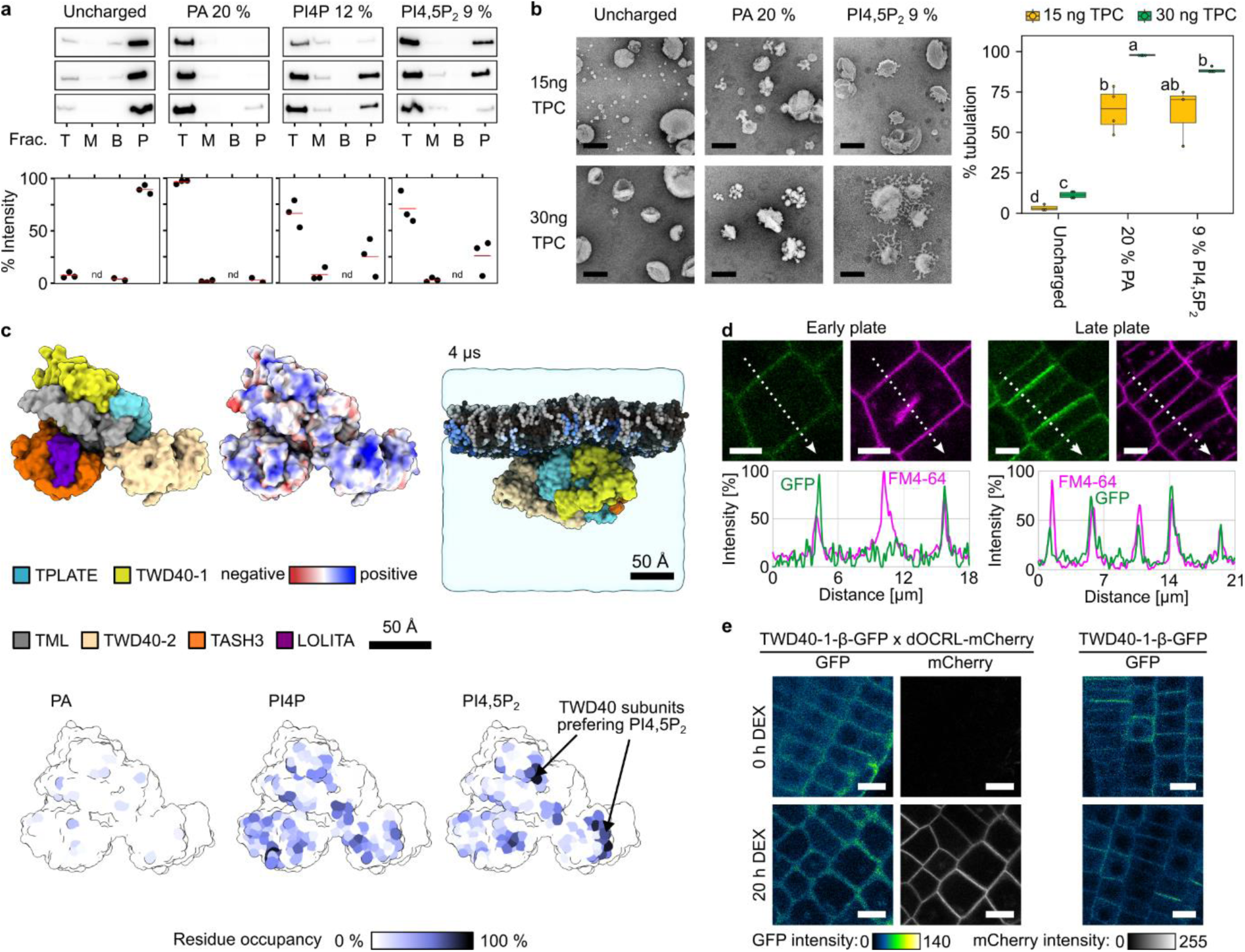
TPC establishes domain-specific lipid interactions. a) Large unilamellar vesicles (LUV)-flotation assay using purified TPC from Arabidopsis PSB-D suspension cells and corresponding quantification of the three independent replicas are shown. Samples following density centrifugation were divided into top fraction (T), middle fraction (M) bottom fraction (B) and pellet (P), frac. = fraction. TPC remains in the pellet fraction (P) in LUVs consisting of uncharged lipids, indicating absence of lipid binding. LUVs containing 20 % of phosphatidic acid (PA) retain TPC in the top fraction (T), while TPC remains detectable in the pellet (P). Similar results were achieved with LUVs containing 12 % phosphatidylinositol 4-phosphate (PI4P) and 9 % phosphatidylinositol 4,5-bisphosphate (PI4,5P_2_). TPC interacts strongest with LUVs containing PA since the complex is almost exclusively detectable in the top fraction (T). nd = not detectable. b) LUV tubulation assay using purified TPC from Arabidopsis suspension cells, visualized by negative stain electron microscopy and quantification of the amount of tubulation observed. TPC tubulates LUVs in a charge and concentration dependent manner. The scale bars equal 200 nm. Letters denote statistically different classes at the 0.01 level. c) Molecular dynamics (MD) simulations revealed the lipid binding sites of the individual TPC domains. The truncated TPC hexamer was used to perform the simulations. The individual subunits are colored according to the legend (top left panel). The top middle panel visualizes the electrostatic potential mapped on the TPC hexamer. The top right panel shows the complex associated with a proxy of the plasma membrane in one of one MD replicas (see methods). Phosphatidylcholine is depicted in black, phosphatidylethanolamine is depicted in dark grey, PS, PA, PI4P and PI4,5P_2_ are depicted in shades of blue. The three panels below visualize the residues on the membrane binding interface of the TPC hexamer that interacted with the selected anionic phospholipids (PA, PI4P, PI4,5P_2_). The scale bars equal 50 Å. d) Representative confocal images showing the localization of the TWD40-1-β-GFP construct, counterstained by the styryl dye FM4-64. TWD40-1-β-GFP is recruited to the PM and the late cell plates but absent from the early ones. The graphs visualize the intensity of both channels along the dotted arrow that is depicted in the images. Scale bars equal 5μm. e) Representative images of Arabidopsis epidermal root cells expressing TWD40-1-β-GFP, combined with the iDePP tool, which inducibly allows to reduce PI4P levels at the PM. Upon induction of the phosphatase (MAP-mCherry-dOCRL) by dexamethasone (DEX), the PM recruitment of TWD40-1-β-GFP is abolished. Dexamethasone treatment alone does not affect PM recruitment of TWD40-1-β-GFP. Scale bars equal 10 μm.

A C-terminal GFP fusion construct of the two N-terminal β-propellers of TWD40-1 was expressed *in planta* (Extended Data Figure 8 a). The construct displayed PM localization together with a strong signal at late cell plates, while being absent at early cell plates (Figure 3 d). This subcellular localization matches the occurrence of PI4,5P_2_^32–34^. To confirm PI4,5P_2_ specificity, we inducibly depleted the PM of PI4,5P_2_ via the iDePP system^33^. Induction of the phosphatase removed the TWD40-1 β-propeller construct from the PM, confirming that the β-propellers bind PI4,5P_2_.

To summarize, the MD simulations corroborated by the data *in planta* together with the charge distribution of the TPC structural model clearly identified a PM binding interface of the TPC with domain-specific lipid preferences (Figure 3). The domain specificities for particular lipids likely reflect the different roles of the TPC throughout the process of endocytosis, or might provide robustness to the process of CME *in planta*.

### TPC functionality requires flexibility

The 210 residue long disordered linker of TML is conserved throughout the plant kingdom. It connects the longin domain with the μHD (Extended Data Figure 5 c). Interestingly, the AP-2 counterpart of TML, AP-2μ, which positions its μHD close to the membrane, also features a disordered linker connecting the two domains. This linker is considerably smaller in AP-2μ than it is in TML (Extended Data Figure 5).

To address the importance of the linker, we replaced the long linker of TML with the shorter AP-2μ linker and generated the construct TML-AP-2μ. A C-terminal GFP fusion of TML-AP-2μ was tested for its functionality via its ability to complement the pollen lethal phenotype of *tml-1* (Figure 4 a). Expression of TML-AP-2μ-GFP was similar to TML-GFP control lines (Extended Data Figure 8 d). Heterozygous *tml-1* plants produce WT offspring plants and heterozygous offspring plants carrying the *tml-1* mutation in an equal ratio^3^. Expression of functional constructs increases the *tml-1* T-DNA transfer ratio (Figure 4 a). Expressing TML-AP-2μ-GFP did not produce more than 50 % of offspring plants carrying the T-DNA mutation. This shows that TML-AP-2μ is not functional, in contrast to the TML-GFP construct containing the long linker (Figure 4 a & b).

**Figure 4.**
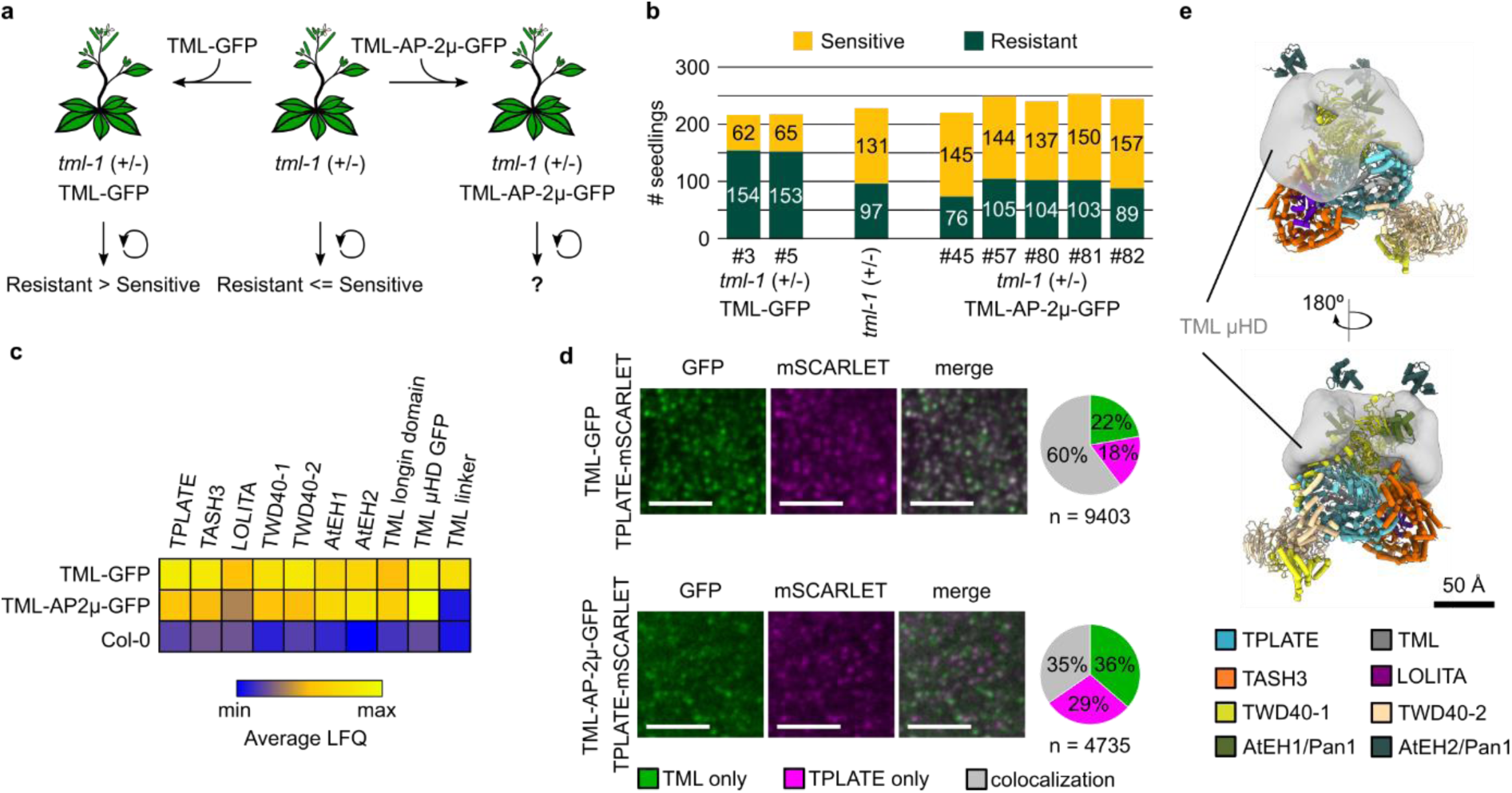
TPC function requires structural flexibility of the TML uHD. a) Schematic representation of the complementation assay of the male sterility phenotype of the *tml-1* mutant. Heterozygous *tml-1* plants generate an equal amount of wild type and heterozygous mutant offspring plants due to the male sterility phenotype of the mutation. Plants expressing functional TML proteins complement the male sterility phenotype and thereby increase the transfer of the T-DNA (visualized by the observed number of resistant offspring) to the next generation. b) The graph shows that the chimeric TML-AP-2μ construct is not functional as the number of seedlings sensitive to Sulfadiazine (wild type) does not exceed the amount of resistant (*tml-1* heterozygous) offspring plants, similar to the non-transformed *tml-1* mutant plants. In contrast, TML-GFP expressing plants are functional as they produce an increased proportion of resistant plants. c) Heat map representation of mass-spectrometry results of the co-immunoprecipitation assay with anti-GFP, comparing TPC formation containing TML-GFP or TML-AP-2μ-GFP in the Col-0 background. Both constructs can still enrich other TPC subunits, indicating that the causality of the non-functionality of TML-AP-2μ-GFP does not lie within impaired assembly of TPC. LFQ = label-free quantification. d) Representative images showing the co-localization of TPLATE-mSCARLET with either TML-GFP (top panels) or TML-AP-2μ-GFP (bottom panels). The pie chart quantifies the colocalization observed. Gray = colocalization, magenta = TPLATE only and green is TML only. To control for random associations of fluorescent signals at the PM, the GFP channel was rotated by 90° to the right and then re-analyzed (Extended Data Figure 8 e). Scale bars equal 5 μm. e) Structural representation of the TPC model with indication of the volume that the μHD domain of TML can occupy (gray). The colors of the TPC subunits are according to the legend. The scale bar equals 50 Å.

As TML-AP-2μ cannot functionally complement the lack of TML, we switched to a Col-0 background to investigate how the TML linker affects TPC formation. In GFP-trap MS experiments, all major TPC subunits were significantly enriched by both constructs (Figure 4 c). The LOLITA subunit was underrepresented in the TML-AP2μ-GFP experiments, likely because of its small size that makes it more difficult to detect. The TML-AP-2μ-GFP bait, however, showed weaker interactions with the other complex subunits compared to the control, indicating a destabilization of TPC, similar to what we observed for the WDXM2 isoform of TPLATE^35^ (Figure 4 c). To visualize TPC formation *in planta*, we introduced TPLATE-mSCARLET into our TML-GFP and TML-AP-2μ-GFP lines. While TML-GFP largely co-localized with TPLATE-mSCARLET, this was not the case for TML-AP-2μ-GFP (Figure 4 d) or for the rotated control images (Extended Data Figure 8 e). The discrete signals in the red and the green channel are likely attributed to either TPC containing endogenous TML or to monomeric TML membrane association, as observed before^16^ (Figure 4 d).

Our combined imaging and proteomics results agree with the formation of an octameric TPC containing TML-AP-2μ-GFP in the cytosol. The flexible linker is not required for membrane recruitment of TML as a single subunit, in agreement with previous results^6^. However, it appears essential for stable PM recruitment of TPC. This would explain the reduced co-localization with other subunits at the PM. Unstable membrane recruitment is likely causal to the non-functionality of TML-AP-2μ-GFP. This is in accordance with our integrative model that visualizes the possible space the μHD of TML can occupy because of its long linker (Figure 4 e).

### Structured domains deform the membrane

TPC is recruited to the PM as an octamer^16^. The AtEH/Pan1 subunits remain at the PM when the TPLATE subunit of TPC is destabilized by heat^35^ and this correlates with the formation of flat clathrin lattices instead of curved clathrin coated pits at the PM in unroofed root protoplasts^8^. Lipid-nucleated phase-separation at the PM of the AtEH/Pan1 TPC subunits serves as a dynamic recruitment platform for early and late endocytic players including clathrin^7^. The above observations imply that AtEH/Pan1-dependent condensate formation does not suffice for initial membrane deformation. We therefore analyzed the role of the structured TPC domains in membrane deformation.

Our MD simulations using the hexamer model showed that, in 3 out of 5 simulation repeats, the TPC hexamer generated membrane curvature (Extended Data Figure 7 e). To computationally test whether TPC-mediated induction of membrane curvature can lead to membrane tubulation, we utilized extensive coarse-grained MD simulations with an asymmetric membrane^36^. Prolonged membrane binding of a single TPC complex to a complex asymmetric membrane, nor the asymmetric membrane alone never resulted in the formation of a membrane tubule (Extended Data Figure 9 a-d). Given that the AtEH/Pan1 subunits have affinity for themselves^7^ and that TPC has several domains capable of lipid binding independent of the AtEH/Pan1 subunits, we assumed that TPC complexes would arrange themselves circular around a membrane-nucleated condensate formed by the AtEH subunits (Figure 5 a-c). This would be in agreement with the model that positions TPC at the edge of a forming clathrin coated pit^2,8^. Therefore, we placed 3 hexamers of TPC close to each other on a complex asymmetric membrane (Figure 5 c). Using this setup as a starting point, 4 out of 5 coarse-grained MD simulations ended in successful membrane deformation, with the TPC molecules residing at the base of the forming bottleneck (Figure 5 d-e, Extended Data Figure 9 e).

**Figure 5.**
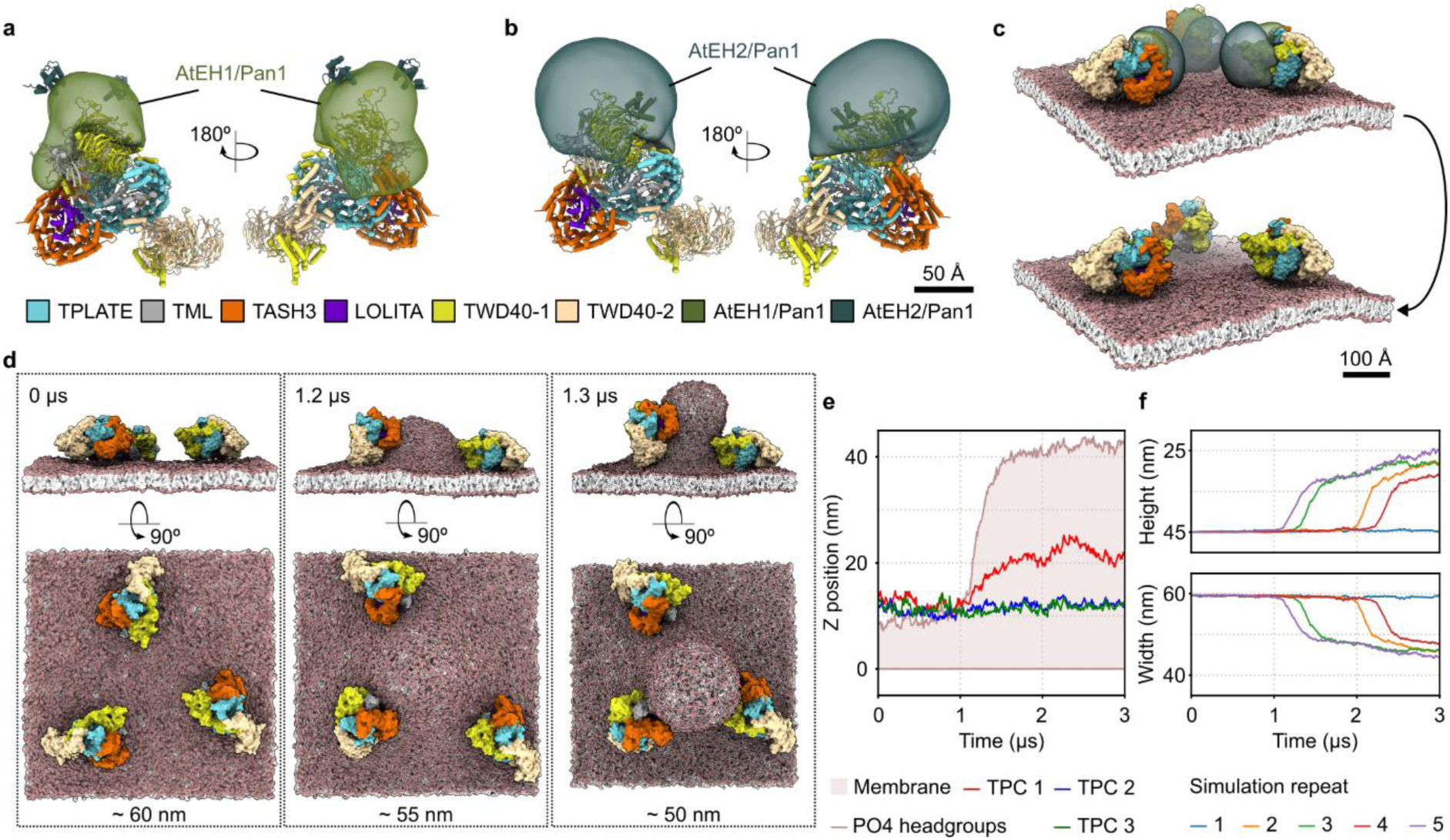
The structured domains of TPC provide the force for membrane deformation. a) and b) TPC structure with indication of the volume that can be occupied by AtEH1 and AtEH2. The scale bar equals 50 Å. c) Positioning of three TPCs for molecular dynamics (MD) simulations. Due to the phase separating capacities of the AtEH subunits, the TPCs were positioned on the membrane with the volumes of the AtEH subunits facing each other (upper panel). Due to their intrinsic disorder, the AtEH subunits were omitted from the MD simulations after the complexes were positioned on the membrane (lower panel). The scale bar equals 100 Å. d) Side and top view images of representative timepoints from the MD simulation. The 0 µs time point represents the starting point and the membrane is roughly 60 nm x 60 nm in size. At 1.2 µs, the membrane starts to deform., leading to a reduction in the membrane size to roughly 55 nm x 55 nm. At 1.3 µs, the membrane deformation proceeds and the size of the membrane is further reduced to roughly 50 nm x 50 nm. e) Visualization of the membrane deformation along the Z axis (brown line and area) over the time of one selected simulation repeat, alongside with the positions of the center of mass of the three TPCs (red, blue and green lines). f) Visualization of the deformation of the simulation box over the time of the simulation. If the membrane deformation occurs (4 out of 5 simulation repeats), the simulation box reduces in width and increases in height. Different colors indicate different simulation repeats.

In conclusion, our model together with previously published data elaborates how the disordered parts of TPC play a crucial role in the recruitment of the complex to the PM (Figure 4 d & e)^7^, while the structured scaffold of TPC with its curved PM binding site is responsible for membrane deformation and tubule generation (Figure 3 c, Figure 5 d-f). Our model for plant CME would therefore position AtEH/Pan1-dependent condensate formation as a nucleation and recruitment module, orchestrating the association of the various players, including AP-2 and clathrin to the endocytic site. The growing pit will likely deplace TPC to the edge of the clathrin-coated pit, similar to the situation in animals with FCHo and EPS15^37^. The highly flexible/disordered proteins FCHo, intersection and EPS15 in animals, or their counterparts Sla1p, Syp1p, Ede1p and End3p in yeast, all share functional domains with TPC that are conserved in their role during CME execution^2^. These players, however, lack the structured scaffold of an adaptor protein complex and vesicle formation occurs in concert with actin-dependent force generation^2,38^. Our results demonstrate that the structured scaffold of TPC is sufficient for membrane deformation, which provides an answer to the long-standing question of why plants do not require actin to drive CME^39^.

## Methods

A detailed description of the methods can be found in the Extended Methods.

### Cloning/molecular work

Genes were amplified via polymerase chain reaction (PCR) and then cloned via either Gibson assembly, Golden Gate cloning, or via Gateway BP and LR recombinations. An extensive list of primers and plasmids that were generated can be found in the Extended Data Table 1.

### PSB-D Cell culture work

To purify TPC from Arabidopsis PSB-D cell suspension cultures, Dark-grown cultures were transformed with recombinant TPLATE containing a C-terminal affinity tag consisting of a TEV cleavage site, GFP and a StrepTagII tag. The transformation and maintenance of the culture was done as described^40^.

### TPC Purification from PSB-D cultures

200 g – 600 g PSB-D frozen culture pellets were ground to a very fine powder in liquid nitrogen with a Braun MultiQuick 7 blender, and subsequently with pestle and mortar. The powder was dissolved and thawed in a lysis buffer (100 mM Tris/HCl pH 8, 150 mM NaCl, 0.5 mM TCEP, 1 tablet Roche Protease inhibitor/50 mL solution) using 1 ml of buffer per g of powder. Benzonase was added to a final concentration of 1 : 10 000 and the solution was incubated on ice for 30 minutes. Afterwards, debris was removed via two subsequent centrifugations. 1 mL of a 50 % suspension of StrepTactin XT 4flow HC beads was equilibrated, by washing it 4 times with lysis buffer. The equilibrated beads (0.5 mL bead volume) were added to the cleared lysate and then incubated for 3 h – 4 h at 4 °C while slowly rotating. The beads were washed with the wash buffer at least six times with nine bead volumes (= 4.5 mL) per step. To elute TPC, one bead volume (= 0.5 mL) of wash buffer containing 0.1 mg/ml TEV was added and the beads were incubated rotating overnight at 4 °C. Equilibrated Ni Sepharose 6 Fast Flow beads were added to the sample from the TEV elution and rotated for 2 h at 4 °C. After incubation, the beads were harvested at 50 x g at 4 °C and washed three times with the wash buffer. Depending on the subsequent in vitro analysis the buffer was changed to accommodate for downstream experiments.

### SDS page analysis and Western blot

Samples were boiled for 8 minutes at 75 °C. The boiled samples were then loaded on a BioRad Mini-PROTEAN® 4-20 % stain-free gradient TGX gel. Stain-free gels were developed using a BioRad ChemiDoc imaging system or the protein sample was transferred to a 0.2 μm PVDF membrane (BioRad) via the Trans-Blot turbo transfer system from BioRad. Afterwards the blots were developed with antibodies listed in the source file.

### GFP trap MS experiments

The Miltenyi Biotec μMACS GFP isolation kit standard protocol for GFP trap experiments was used with adapted buffers to make sure that TPC remained stable. LC-MS/MS analysis was done on an Ultimate 3000 RSLCnano system in-line connected to a Q Exactive mass spectrometer (Thermo). The Thermo raw files were processed using the MaxQuant software (version 2.2.0.0)^41^. The MaxQuant protein groups result file was uploaded in Perseus (version 1.6.15.0)^42^ for Label Free Quantification (LFQ) based differential analysis. LFQ intensity values were log2 transformed. Identifications were filtered for minimal three valid values in at least one group. Missing values were imputed with values around the detection limit, randomly drawn from a normal distribution with a width equal to 0.3 and a downshift equal to 1.8. Two-sided tests were performed, and significantly enriched proteins were determined by permutation based FDR calculation, using thresholds FDR = 0.02 and S0 = 1. The results, including the imputed dataset, can be found in Extended Data Table 2.

### XL-MS analysis

Cross-linking reactions were performed in solution with 13.5 μg TPC per reaction. 1.2 mM DSSO (Thermo Fisher A33545) was added, the reaction was incubated at room temperature for 1 h and terminated by adding 180 μL of 50 mM NH_4_HCO_3_ pH 8.0. To prepare the samples for MS, they were reduced with 5 mM DTT for 40 minutes at room temperature. The reduced samples were alkylated 30 minutes at room temperature with 15 mM Iodoacetamide. The samples were digested with 1.25 μg Trypsin/LysC at 37 °C overnight and for another 2 hours the next morning by adding fresh Trypsin/LysC solution. The trypsinated peptides were dried via speedvac and then analyzed via LC-MS/MS.

Samples were injected for LC-MS/MS analysis on an Ultimate 3000 RSLCnano system in-line connected to an Orbitrap Fusion Lumos mass spectrometer (Thermo). Thermo raw files were analyzed using MS Annika 2.0^43^ integrated in Proteome Discoverer 2.4.1.15. For Merox 2.0^44^ analysis, raw files were first converted to mzML using MSConvertGUI v3.0.21084. Data analysis was performed against a database containing the 8 subunits of the TPLATE complex. The BS3 dataset from our previous publication^6^ was re-analyzed. The software specific settings and identified cross-links can be found in Extended Data Table 3.

### Single band SDS page MS analysis

The SDS page gels were stained with Colloidal Coomassie to visualize and excise the bands. Excised bands were destained every 30 minutes with HPLC grade water. Samples were reduced by replacing the water with a reducing buffer (50 mM NH_4_HCO_3_ pH 8, 5 mM DTT) and incubated for 40 minutes at room temperature while shaking. The reducing buffer was changed for an alkylation buffer (50 mM NH_4_HCO_3_ pH 8, 50 mM Iodoacetamide) and the samples were incubated for 30 minutes at room temperature. After alkylation the samples were washed two times with water before continuing with the in-gel trypsin digestion. To release all digested peptides from the gel plugs, they were sonicated for 5 minutes. The supernatant with the trypsinated peptides was transferred to a clean tube and the plugs were washed with 50 μL replacement buffer (50 mM NH_4_HCO_3_ pH 8, 10 % acetonitrile). The wash fraction was added to the trypsinated peptides and the peptides were then purified via Agilent C18 Omix tips (catalog number: A57003100) as described by the manufacturer.

LC-MS/MS and MaxQuant analysis were performed as described for the GFP trap experiments, with exception of the database used in the MaxQuant search. Here, Araport11plus was used. The results are listed in Extended Data Table 4.

### Polyclonal antibody generation

Peptide-based antibodies against TASH3 (C+LEKVGDVPHKRKKGVFGTK) and LOLITA (KLGADNLKGVKNEEL+C) were developed with BIOTEM. The SDS pages and Western blots confirming the antibody specificity and determining their background signal are depicted in Extended Data Figure 10.

### Design of the TML-AP-2μ chimera

The TML-AP-2μ chimera construct was generated using fragments whose boundaries were defined via multiple sequence alignment. The longin and μHD domain were PCR amplified and the linker sequence was ordered using primers (Extended Data Table 1). All fragments were combined via the GoldenGate cloning system.

### Plant line generation

The TMLp::TML-GFP, TMLp::TML-AP-2μ-GFP and H3.3p::TWD40-1-β-GFP plant lines were generated by floral dip^45^ of Col-0 plants or *tml-1* mutant plants^3^.We selected at least two independent lines for each construct and confirmed the expression level of the introduced heterologous gene via Western blot. For the colocalization analysis of the TML constructs with LAT52p::TPLATE-mSCARLET, the established homozygous TMLp::TML-GFP and TMLp::TML-AP-2μ-GFP expressing lines in the Col-0 background were transformed with a functional LAT52p::TPLATE-mSCARLET construct^46^. To generate the plant lines used for the inducible depletion of PI4,5P2, dOCRL lines^33^ were transformed with the H3.3p::TWD40-1-β-GFP construct. Expression of the TWD40-1-β-GFP construct severely suffers from silencing in between generations, which prevents the establishment of stable homozygous lines. T1 transformants were therefore imaged. An overview of plant lines can be found in Extended Data Table 1.

### Functionality test for TML constructs

The TMLp::TML-GFP and TMLp::TML-AP-2μ-GFP constructs were introduced in the *tml-1* mutant background and the transfer rate of the *tml-1* T-DNA insertion in the T2 generation was evaluated via the Sulfadiazine resistance located on the *tml-1* T-DNA^3^.

### LUV flotation

Lipids (Avanti Polar Lipids) were dissolved in chloroform (PC, PE, and PA) or in chloroform, methanol, and water (PI4P and PI4,5P_2_). Lipids were mixed in ratios 80:20 for PC-PE, 60:20:20 for PC-PE-PA, 68:20:12 for PC-PE-PI4P, and 71:20:9 for PC-PE-PI4,5P_2_ and speedvac dried. The pellets were redissolved in 20 mM HEPES pH 7.5, 250 mM NaCl, 0.5 mM TCEP buffer and incubated for 90 minutes on a rotational wheel. Following the incubation, lipid suspensions were sonicated for 15 minutes in an ultrasonic bath. To obtain unilamellar liposomes of a homogenous size, the lipid suspensions were extruded through a polycarbonate membrane with a 0.2 μm pore size (Whatman). For the co-flotation assays, 150 nmol liposomes per sample were mixed with purified TPC and incubated at room temperature for 45 minutes. After incubation, 100 μl of 60 % sucrose (w/v) was added to the sample resulting in a concentration of 30 % sucrose (w/v). The mixtures were transferred to centrifugation tubes, overlaid with 250 μl 25 % sucrose w/v and 50 μl 20 mM HEPES pH 7.5, 250 mM NaCl, 0.5 mM TCEP buffer and centrifuged for 2 hrs at 175,000g using S120-AT2 fixed angle rotor (Sorvall), 22 °C.

### LUV tubulation

Lipids were treated identical as for the flotation assays. Pellets were redissolved in 500 μl 20 mM HEPES, 15 % (w/v) raffinose, 0.5 mM TCEP and incubated for 90 minutes on the rotational wheel. lipid suspensions were sonicated and extruded. Liposomes were diluted up to 1 ml with 20 mM HEPES, 250 mM NaCl, 0.5 mM TCEP buffer, pH 7.5 and centrifuged 30 min at 74000g using S120-AT2 fixed angle rotor (Sorvall), 22 °C. The pellet was resuspended in 500 μl buffer and the centrifugation step was repeated. The supernatant was discarded, and the pellet was resuspended in 100 μl buffer. For the tubulation reaction, liposomes were diluted to a final concentration of approximately 2 mM. 10 μL of freshly purified TPC in tubulation buffer (20 mM HEPES/NaOH pH 7.6, 250 mM NaCl, 50 mM Maltose, 0.5 mM TCEP) was gently mixed with 20 μL of liposomes and incubated for 45 minutes on a rotating wheel at 6 rpm. For the no protein control, 10 μL of tubulation buffer was added to 20 μL of liposomes. Afterwards the samples were diluted 4 x in the tubulation buffer and used right away for negative stain EM grid preparation. Micrographs were imaged on a JEOL 1400+ microscope with a LaB6 filament at 120 kV acceleration voltage using a TVIPS F416 CCD camera at 25 000 x magnification.

### Confocal microscopy

Colocalization analysis between TPLATE and TML was performed on a Nikon Ti microscope with the Perfect Focus System (PFSIII) for Z-drift compensation, equipped with an Ultraview spinning-disk system (PerkinElmer) and two 512×512 Hamamatsu ImagEM C9100-13 EMccd cameras. The microscope was controlled via the Volocity V7.0.0 software package. IDePP imaging was performed on the spinning disk and on a Leica DMI8 TCS SP8X confocal microscope with a white light laser and hybrid detectors (HyD). Imaging of TWD40-1-β-GFP in combination with FM4-64 staining was performed on the Leica SP8X.

### Negative staining of EM grids

4 μL of purified TPC was applied to glow discharged Formvar FCF400-CU-50 EM grids. After 30 seconds the protein was blotted away and the grid was immediately placed on top of an uranyl acetate drop for 10 seconds. Afterwards the uranyl acetate was removed and the grid was placed on a second uranyl acetate drop for 1 second. After blotting away the uranyl acetate the grid was incubated for 1 minute on the third uranyl acetate drop. Finally, the uranyl acetate was blotted away, leaving a thin film and the grids were dried at room temperature before imaging.

### TPC negative stain EM dataset acquisition

Negative stain micrographs were imaged on a JEOL 1400+ microscope with a LaB6 filament at 120 kV acceleration voltage using a TVIPS F416 CCD camera at 60 000 x magnification corresponding to a magnified pixel size of 1.94 Å/px. Automated data acquisition was performed via Serial EM v4.0.28^47,48^. The dataset was then acquired in a defocus range of −0.5 μm – 3 μm with a step size of 0.5 μm. Before data processing, micrographs with visible artefacts, like drift, astigmatism and protein aggregation were discarded.

### 3D map generation of negative stain dataset via Cryosparc

The nsEM datasets were evaluated with CryoSparc v4.1.2^49^. CTF estimation was performed via CTFFIND4 v4.1.14^50^. Particles were automatically picked using the pre-trained topaz resnet8 model with 64 units, a downsampling factor of 8 and a radius of 8 with topaz v0.2.5^51^.The final set of particles was selected in three rounds of 2D classification using a maximum resolution of 15 Å, a circular mask diameter of 270 Å (resulting in a box size of 324, equalling roughly 1.5 x the longest axis of TPC) and a white noise model.

### Integrative structure modeling of the TPC

The Python modeling interface of the integrative modeling platform (IMP) package version 2.17^23,52^ was used to build the TPC structure.123 and 126 unique intra- and intermolecular BS^3^ and DSSO cross-links were used to construct the scoring function. In addition to the cross-link datasets, the excluded volume restraints, the sequence connectivity restraints and the GMM map were added to the scoring function. Replica Exchange Gibbs Monte Carlo algorithm^53^ was used to sample structures satisfying input restraints.

### Molecular dynamics simulation

All simulations were performed using the GROMACS software^54^. MARTINI 2.2 coarse-grained (CG) force-field^55^ was used for the simulations of TPC with a membrane. The relaxed TPC structure was converted to coarse-grained (CG) representation using the CHARMM-GUI web server^56,57^. Elastic Network in Dynamics (ElNeDyn) was applied to retain the secondary and tertiary structure^58^.

### Statistical analysis

The R package in R studio was used for statistical analysis (https://www.R-project.org/). For multiple comparison the multcomp package was used^59^.

## Supporting information

video 1 Integrative modeling of the TPC structure

video 2 Structured domains of the TPC deform the membrane

## Data Availability

The nsEM map is deposited in the EMDB database (XXX). All data files related to integrative modeling and molecular dynamics simulations are deposited in the Zenodo repository: XXX. The mass spectrometry data are deposited to the ProteomeXchange Consortium via the PRIDE partner repository with the dataset identifier XXX. Source data are provided with this study.

## Acknowledgements

The authors would like to thank Andreas De Meyer and Jonathan Michael Dragwidge for help with live cell imaging, Frederik Delaere and Frederik Coppens for the access and maintenance of the server and Jonah Nolf for help with Äkta-based protein purification. We would like to thank Francis Impens and An Staes for help with XL-MS and Savvas Savvides, Jan Felix and Kenneth Verstraete for their help with the Cryo plunger and with the CryoSparc software package. We would also like to thank Rouslan Efremov for constructive discussions on the purification and imaging strategies. This work was supported by the European Research Council Grant T-REX 682436 (D.V.D.), the Research Foundation–Flanders (FWO) G017919N (M.K.), The Czech Science Foundation grant nr. 22-35680M (R.P.), project GA UK nr. 154324 (M.N.). Computational resources used for MD simulations were provided by the e-INFRA CZ project (ID:90254), supported by the Ministry of Education, Youth and Sports of the Czech Republic.

## Author contributions

J.M.K., M.N., A.F.C., N.S., M.V., K.Y and E.M. performed experiments. N.D.W and E.V.D.S. generated PSB-D cell cultures. R.P. performed integrative modeling. D.E. performed MS data-analysis. M.F. helped with EM data recording and processing. G.D.J., R.P. and D.V.D designed experiments and supervised the work.

## Ethics declaration

The authors declare no conflict of interest.

## Additional information

### Extended Data Figures and Tables

**Extended Data Fig. 1.**
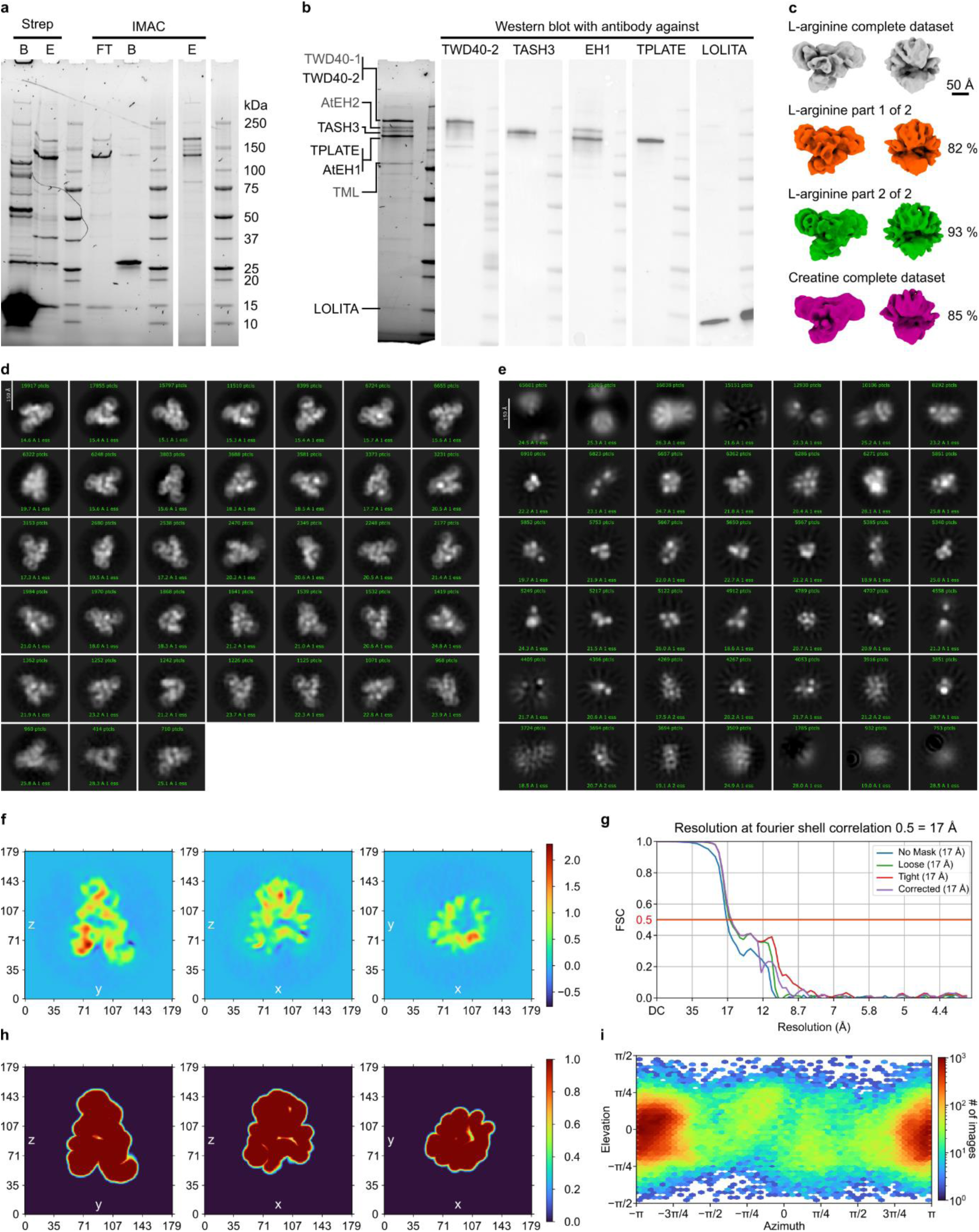
TPC purification and negative stain structure generation. a) Stain free SDS PAGE gels showing the purification progress of TPC from Arabidopsis PSB-D cell suspension culture. The TEV elution during the StrepTag purification elutes TPC well as there is no residual complex on the beads (Strep B). Since the HIS-tag site is not part of the bait, but part of TPC, excessive bait cannot bind the Ni-NTA Sepharose beads (IMAC FT). The established purification conditions allow us to elute TPC selectively while retaining the TEV on the beads (IMAC B & IMAC E), yielding a clean TPC sample. b) To identify the individual subunits, purified TPC was analyzed via Western blots, using antibodies against selected TPC subunits (black font), namely: anti-TWD40-2^60^, anti-TASH3, anti-AtEH1^18^, anti-TPLATE^61^ and anti-LOLITA. As described before^18^ the anti-AtEH1 antibody cross reacts with AtEH2 visible by the second band in the Western blot. Both anti-TASH3 and anti-LOLITA are antibodies developed as part of this research and the specificity is presented in Extended Data Figure 10. The annotation via Western blot was complemented by single band SDS page MS experiments. These results confirmed the Western Blot analysis and allowed the annotation of the remaining subunits (grey font) (Source file). c) The 3D map generated with the full L-arginine dataset (grey) was validated by splitting the dataset into two parts and a 3D map was reconstituted for each part individually (part 1 = orange, part 2 = green). In addition to that, a 3D map was reconstituted from TPC purified in a creatine containing buffer (magenta). These independently calculated maps were then compared via UCSF ChimeraX^62,63^ against the map from the full L-arginine dataset. The correlation value of these pairwise comparisons is depicted next to the corresponding map. As is visible in c), the maps all present the same shape and dimensions. The scale bar measures 50 Å. d)-i) show the 3D reconstruction of the L-arginine dataset. d) After 3 rounds of 2D classification, where particles were discarded, we yielded a final particle set of 156 997 particles. 2D classes corresponding to this final set are depicted in d). e) To exclude that good particles were removed during 2D classification, all discarded particles were re-classified. The resulting 2D classes are presented in e) and no 2D classes of interest were present. f) The densities plotted in the real space slices indicate that TPC particles can be clearly distinguished from the background and noise of the micrographs during 3D refinement. g) The fourier shell correlation (FSC) estimates a 17 Å resolution at 0.5 FSC for the refined 3D map. h) The mask that was determined during the 3D refinement to compute the 3D map is in accordance with the real space slices in panel f) i) The Azimuth graph positions all particles used for the 3D refinement onto the refined 3D TPC map. This results in a plot where one can determine which view angle has more, or less particles. Ideally one would have an even distribution over the whole graph. The uneven distribution visible in this graph indicates a preferred orientation of the TPC on the negative stain EM grid.

**Extended Data Fig. 2.**
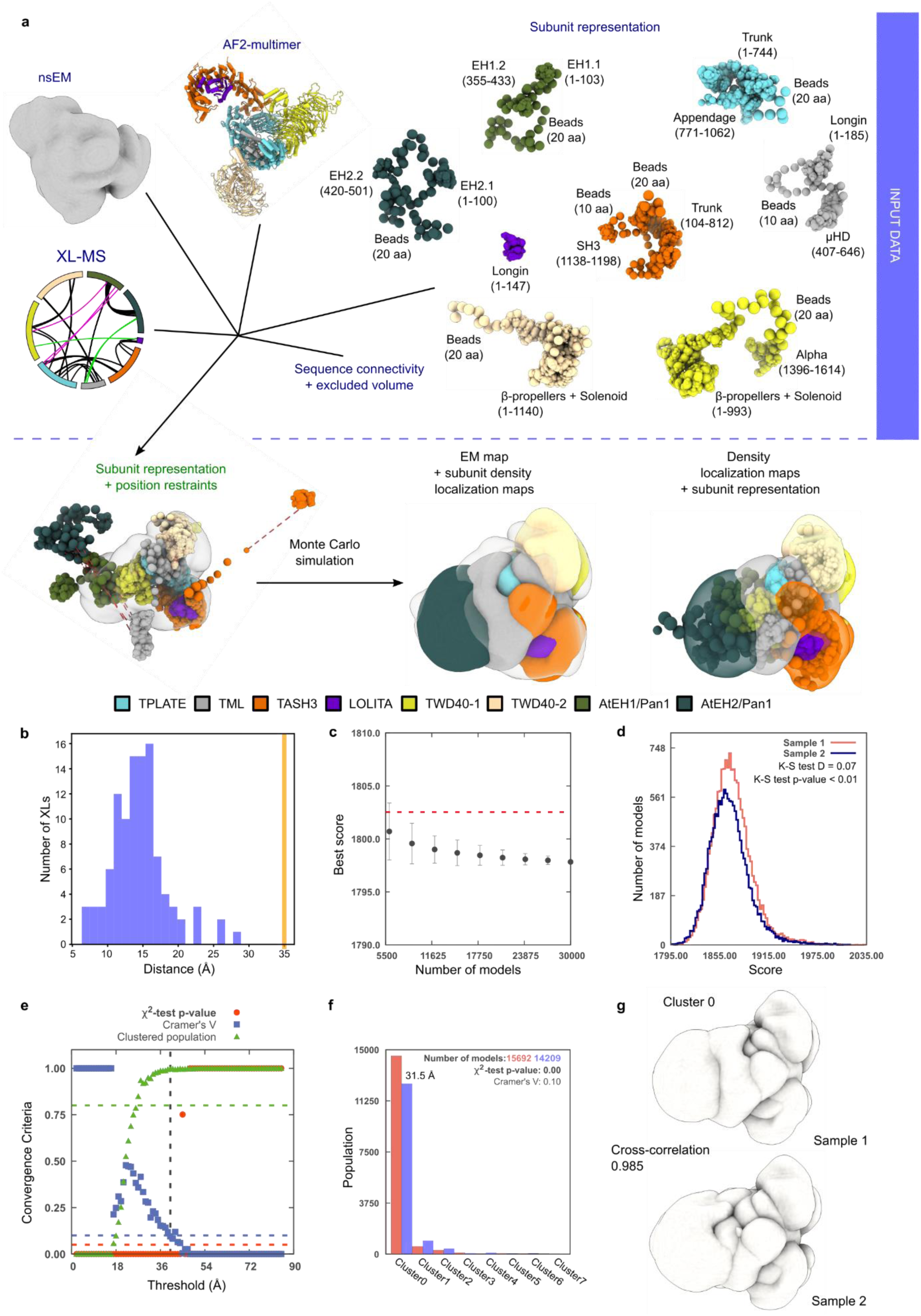
Integrative modeling of TPC and modeling validation. a) The scheme illustrates different stages of the integrative modeling process, including data gathering, representation of the subunits, configurational sampling and model visualization. The input data included the negative stain electron microscopy (nsEM) map, chemical cross-links (XL-MS), multiscale representation of the TPC subunits, TPC subcomplexes modeled by Alphafold2 and stereochemical restraints. b) Distance distribution of obtained chemical cross-links in the top-scoring cluster. The orange line represents the threshold for the consistent cross-links. No cross-link was violated in the top-scoring cluster. c) Convergence of the model score calculated for the ensemble of the good-scoring models. Scoring did not improve after the addition of more independent models. The error bars represent the standard deviation of the best scores, estimated by repeating the sampling 10 times. The red line depicts a lower bound on the total score. d) Splitting of the good-scoring models into two sample populations (blue and light red) resulted in significantly different score distributions (p-value < 0.01), but the magnitude of the difference is, however, negligible (0.07), therefore both score distributions are effectively equal. e) Determining the sampling precision according to^64^. The sampling precision as defined by three criteria; first, the p-value calculated using the χ2-test for homogeneity of proportions (red dots); second, an effect size for the χ2-test is quantified by the Cramer’s V value (blue squares); third, sufficiently large clusters (containing at least 10 models) visualized as green triangles. The vertical dotted gray line indicates the root mean square displacement (RMSD) clustering threshold at which three criteria are satisfied (p-value > 0.05, Cramer’s V < 0.10, and the population of clustered models > 0.80). The calculated sampling precision is 40 Å. f) Using the sampling precision as the threshold, populations of sample 1 (light red) and 2 (blue) form eight clusters. 90.5 % of the models belong to cluster 0, which has a precision of 31.5 Å. g) Localization density maps for Sample 1 and Sample 2 of Cluster 0, visualized here at a threshold equal to one-tenth the maximum. The cross-correlation of the localization density maps of the two samples is 0.985, indicating that the position of TPC subunits in the two samples is identical.

**Extended Data Fig. 3.**
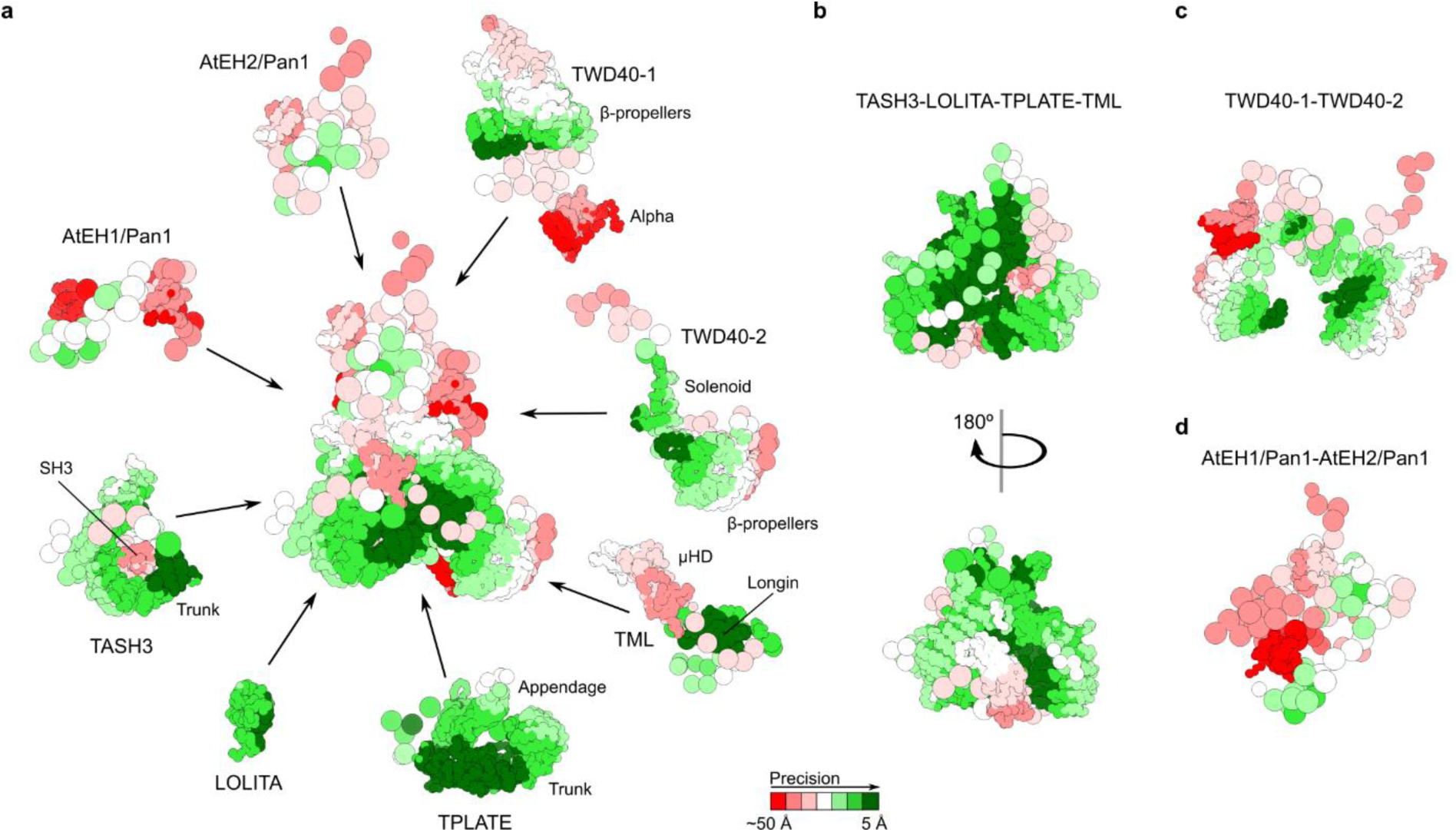
Visualization of the high and low-precision regions of the integrative model. a) Precision of the TPC integrative model as well as that of the individual subunits according to the PrISM method^65^. b) Overall high-precision of the core sub-complex consisting of the TASH3 trunk domain, LOLITA, the TPLATE trunk domain and the TML longin domain. c) Precision of the TWD40 subunits arching the inner core. The structured parts are well resolved with the exception of the all alpha domain (alpha), which contains a flexible linker. d) Precision of the AtEH subunits. As these subunits present the most flexible subunits in the complex, they have the lowest precision in the integrative TPC model.

**Extended Data Fig. 4.**
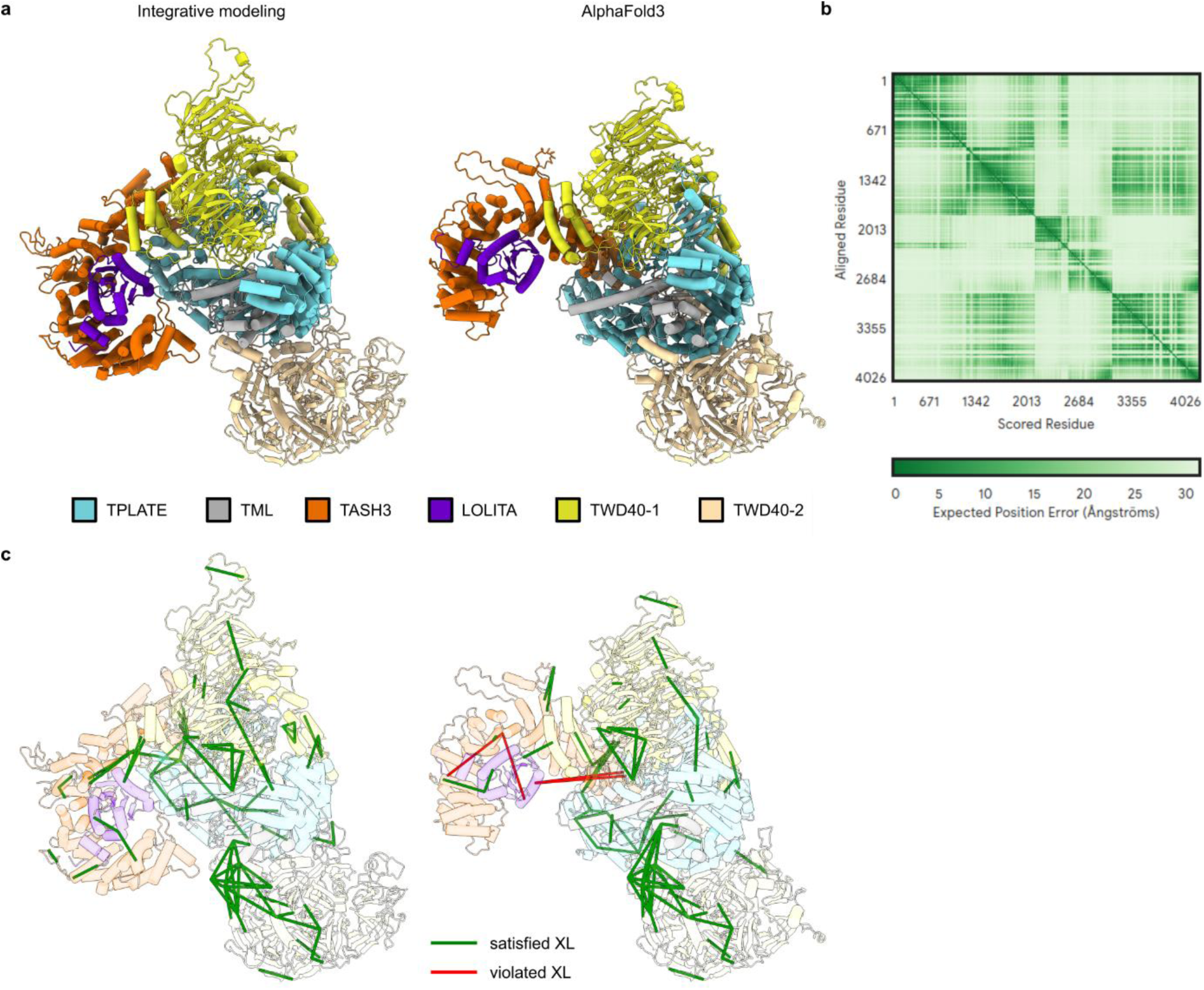
Comparison between the integrative and the AlphaFold3 generated model of the TPC hexamer. a) Ribbon representation of the truncated TPC hexamer obtained by integrative modeling and Alphafold3 (AF3). b) The matrix shows the Predicted Alignment Error (PAE) for the AF3 model of the truncated TPC hexamer. c) Inter- and intramolecular DSSO and BS3 cross-links were mapped on both models. The AlphaFold3 model mainly differs from the integrative structure in the position of the TASH3 trunk domain and LOLITA. The position of the TASH3 trunk domain and LOLITA in the Alphafold3 model is not supported by the identified cross-links.

**Extended Data Fig. 5.**
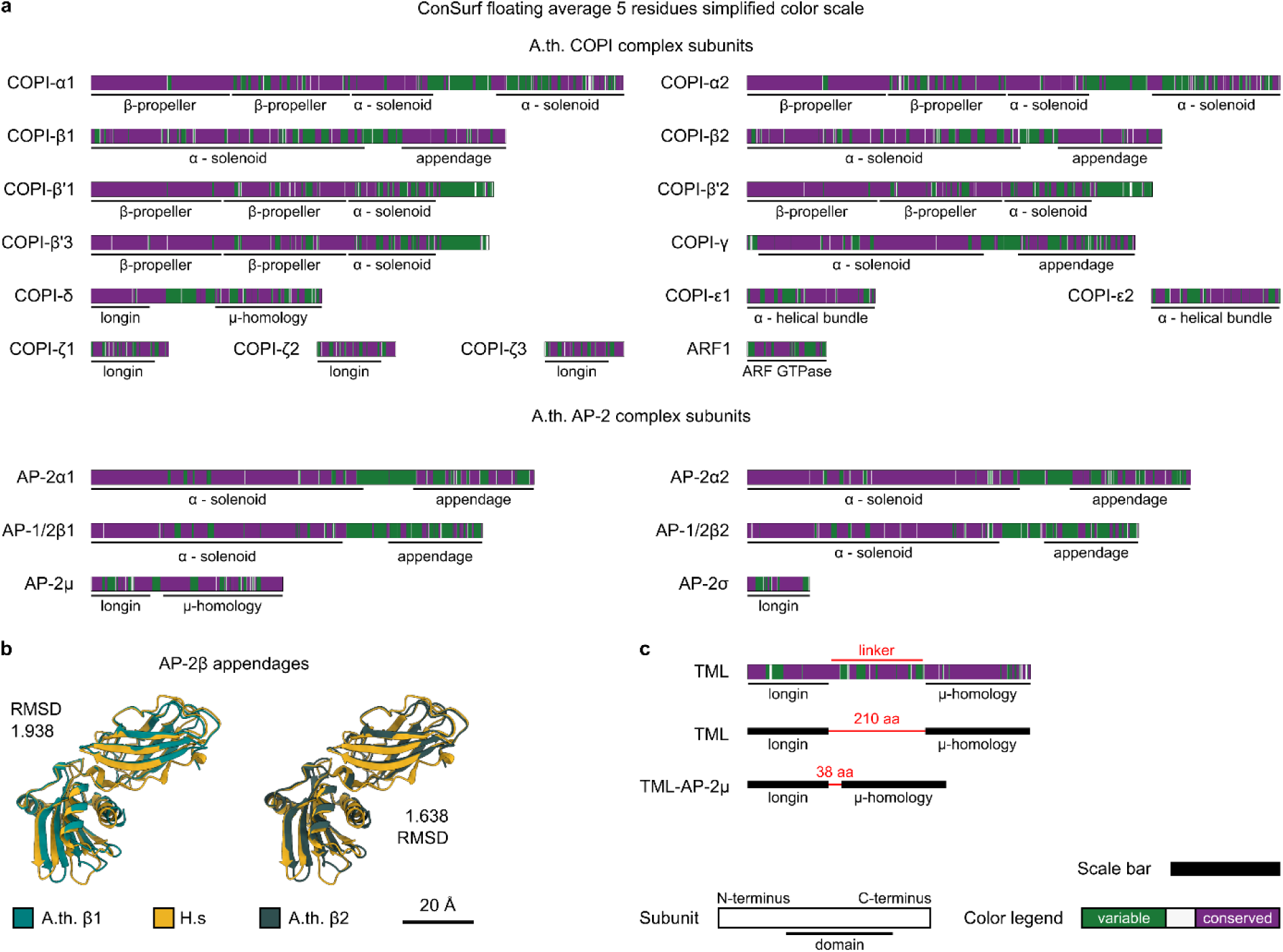
Conservation analysis for structure comparison. a) ConSurf^28^ analysis for all COPI and AP-2 subunits present in *A. th*. A floating average of 5 residues was calculated from the per-residue ConSurf score. This average was then colored green, if the value was below 5 and the sequence therefore variable, or it was colored purple if the score was above 5 and therefore conserved. b) The AP-2β appendages are the least conserved domains in the analysis performed in a). To demonstrate the structural conservation for the AP-2β appendages, structural models of the Arabidopsis AP-2β subunits were taken from the AlphaFold2 database^66^ and the appendage domains were aligned to the published human cryo EM model (PDB 6YAI) with UCSF ChimeraX^62,63^. The AP-2β appendages of both Arabidopsis alleles are structurally nearly identical to their human counterpart, as is evident from their RMSD values. c) To demonstrate the conservation of the disordered TML linker, we performed a ConSurf analysis for TML as in a). This conserved 208 residue linker was replaced with the shorter 38 residue long AP-2μ linker from Arabidopsis in the TML-AP-2μ chimera. The legend below c) applies to a) and c). The scale bar represents 250 residues.

**Extended Data Fig. 6.**
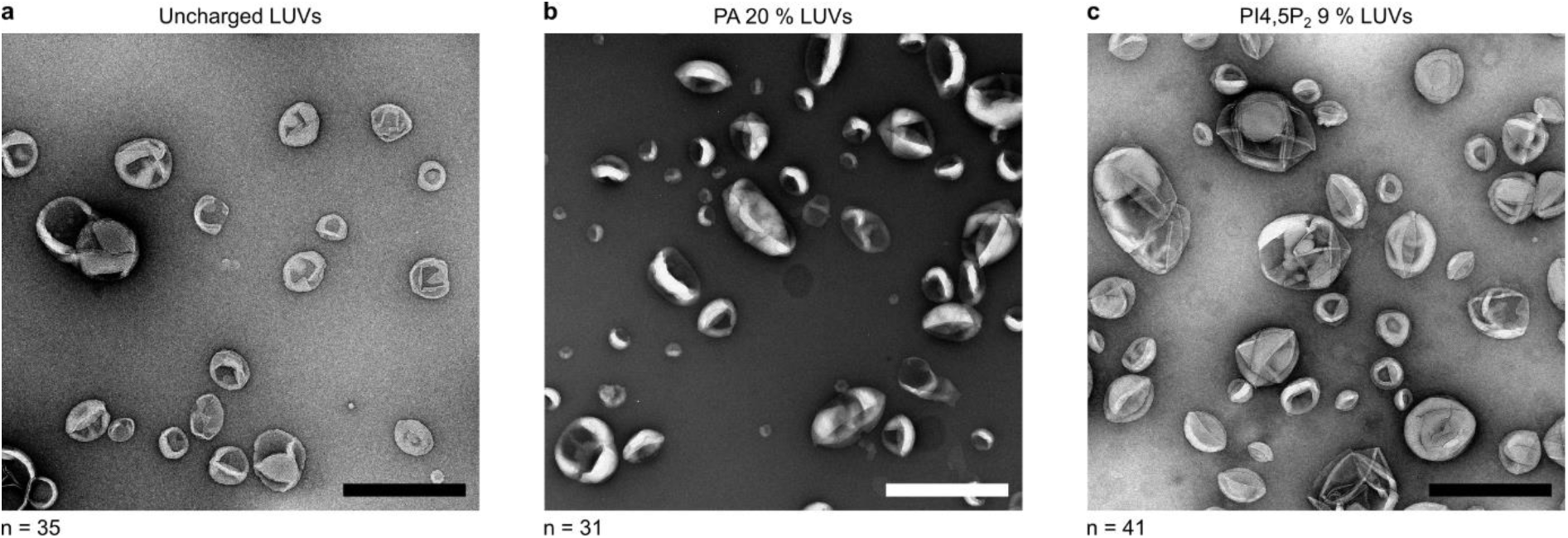
LUVs of different lipid compositions do not tubulate without TPC. a) - c) LUVs with no TPC added do not tubulate. The images are taken at 25 000 times magnification and the scale bars measure 500 nm. a) Depicts uncharged LUVs, while in b) LUVs contain 20 % PA and in c) the PI4,5P_2_ content is 9 %. n = number of acquired micrographs.

**Extended Data Fig. 7.**
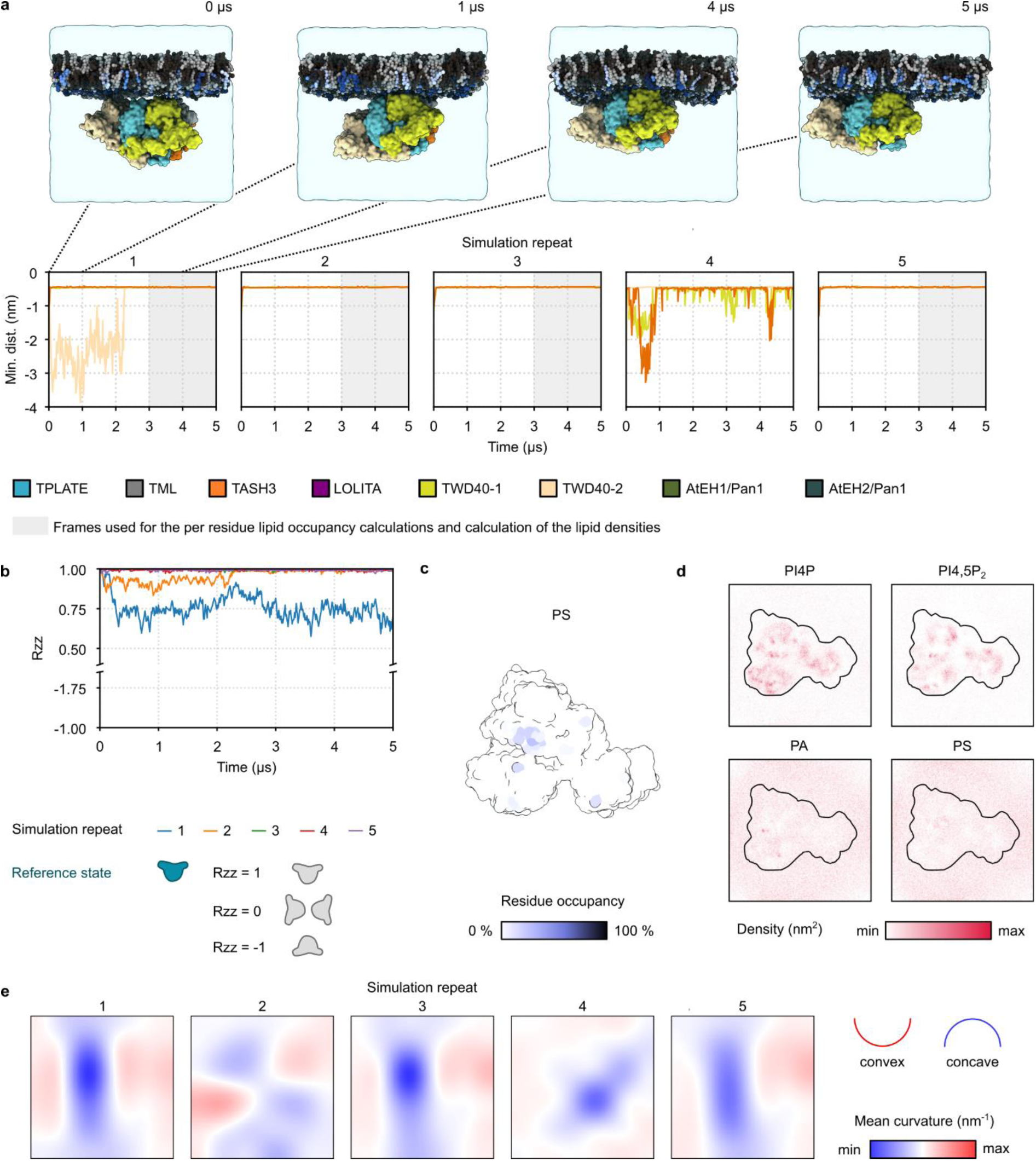
MD simulations reveal TPC membrane interactions. a) Selected snapshots of MD simulation time series showing TPC interaction with the membrane. The minimum distance between TWD40-1, TWD40-2 and TASH3 and the membrane was calculated to monitor TPC-membrane interaction. The last 2 μs of trajectories in which TWD40-1, TWD40-2 and TASH3 simultaneously bind the membrane, were selected for follow up analyses. Phosphatidylcholine is depicted in black, phosphatidylethanolamine is depicted in dark grey, PS, PA, PI4P and PI4,5P_2_ are depicted in shades of blue. b) ZZ component of the rotational matrix visualizing rotation of the TPC in the simulation box in time. 4 out 5 simulation replicas show no rotation of the complex once bound to the membrane. Rzz = 1 no rotation, Rzz = 0 rotation 90°/270°, Rzz = −1 rotation 180° c) Residues of the membrane binding interface of the TPC hexamer interacting with phosphatidylserine (PS). d) PI4P, PI4,5P_2_, PA and PS densities around TPC hexamer. PI4P, PI4,5P_2_ cluster at the membrane binding interface. e) Membrane curvature over the last 2 μs calculated for each MD trajectory. In 3 out of 5 simulation repeats, TPC interacting with the membrane induces the concave membrane curvature.

**Extended Data Fig. 8.**
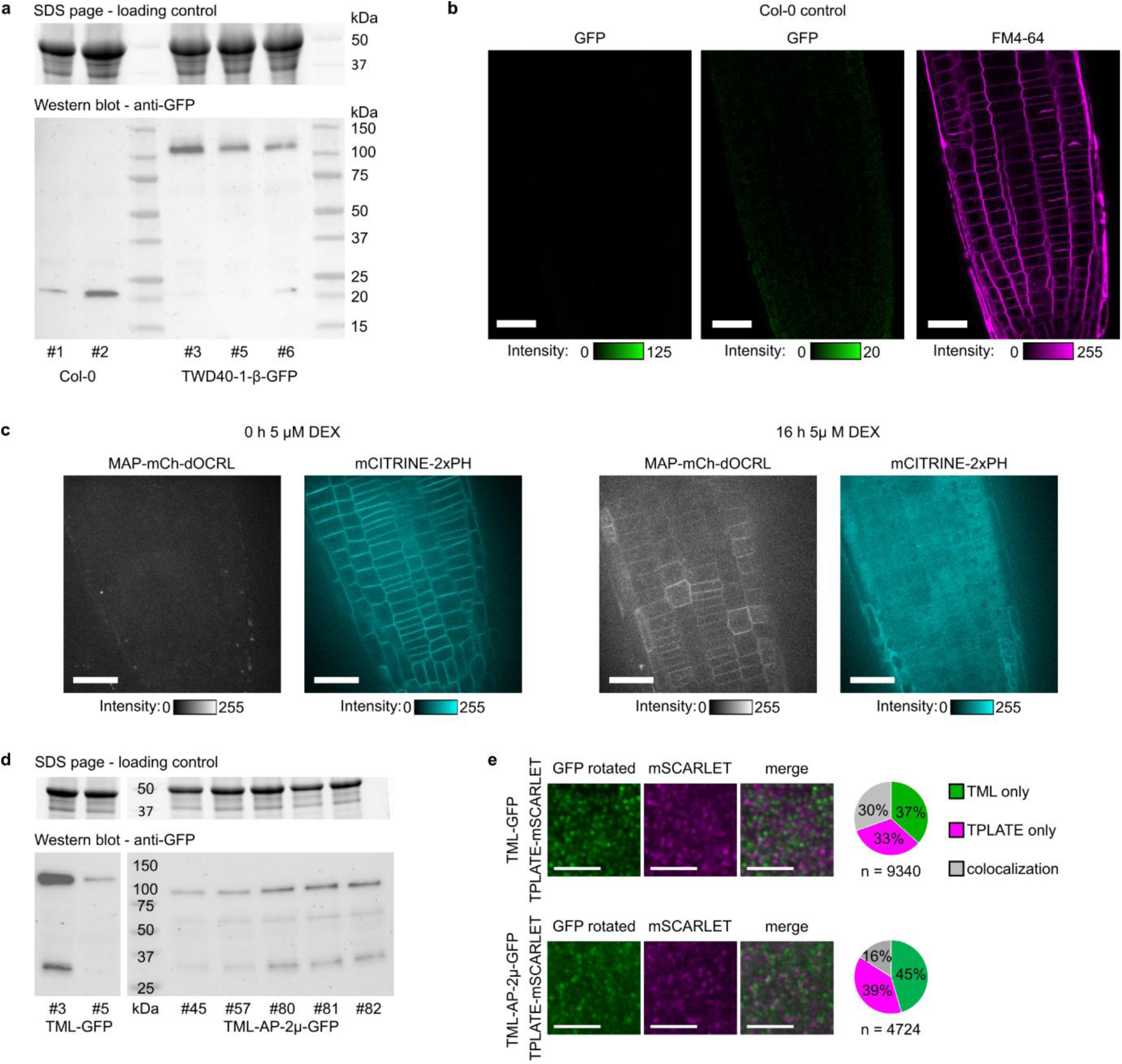
Expression and imaging controls for plant-related experiments. a) Expression analysis of TWD40-1-β-GFP in independent plant lines in Col-0 background that were used for imaging. The stain free gel that serves as loading control is shown on top. The anti-GFP Western blot shows that the constructs are stably expressed. b) Bleed through control images for the data shown in Figure 3 d. The left and middle panels show the signal of a Col-0 root using our imaging conditions at different LUT levels as well as the FM4-64 channel image. Our imaging conditions, using a combination of three excitation lines for GFP does not result in FM4-64 bleed through. The scale bars equal 25 μm. c) Eight-bit full width LUT control images for the iDePP system shown in Figure 3 e. The PI4P PH domain marker (mCITRINE-2xPH) localizes to the PM before dOCRL induction and becomes cytoplasmic after Dexamethasone-dependent induction of dOCRL expression (16 h 5 μM). Scale bars equal 25 μm. d) Expression analysis of independent TML-GFP and TML-AP-2μ-GFP lines in *tml-1* background. The Western blot indicates both constructs are expressed at comparable levels. e) To control for random associations of TML-GFP (top panels) or TML-AP-2μ-GFP (bottom panels) with TPLATE-mSCARLET in the colocalization analysis, the green channel was rotated by 90° to the right and the images were re-analyzed. The pie chart quantifies the colocalization observed. Gray = colocalization, magenta = TPLATE only and green is TML only. Scale bars equal 5 μm.

**Extended Data Fig. 9.**
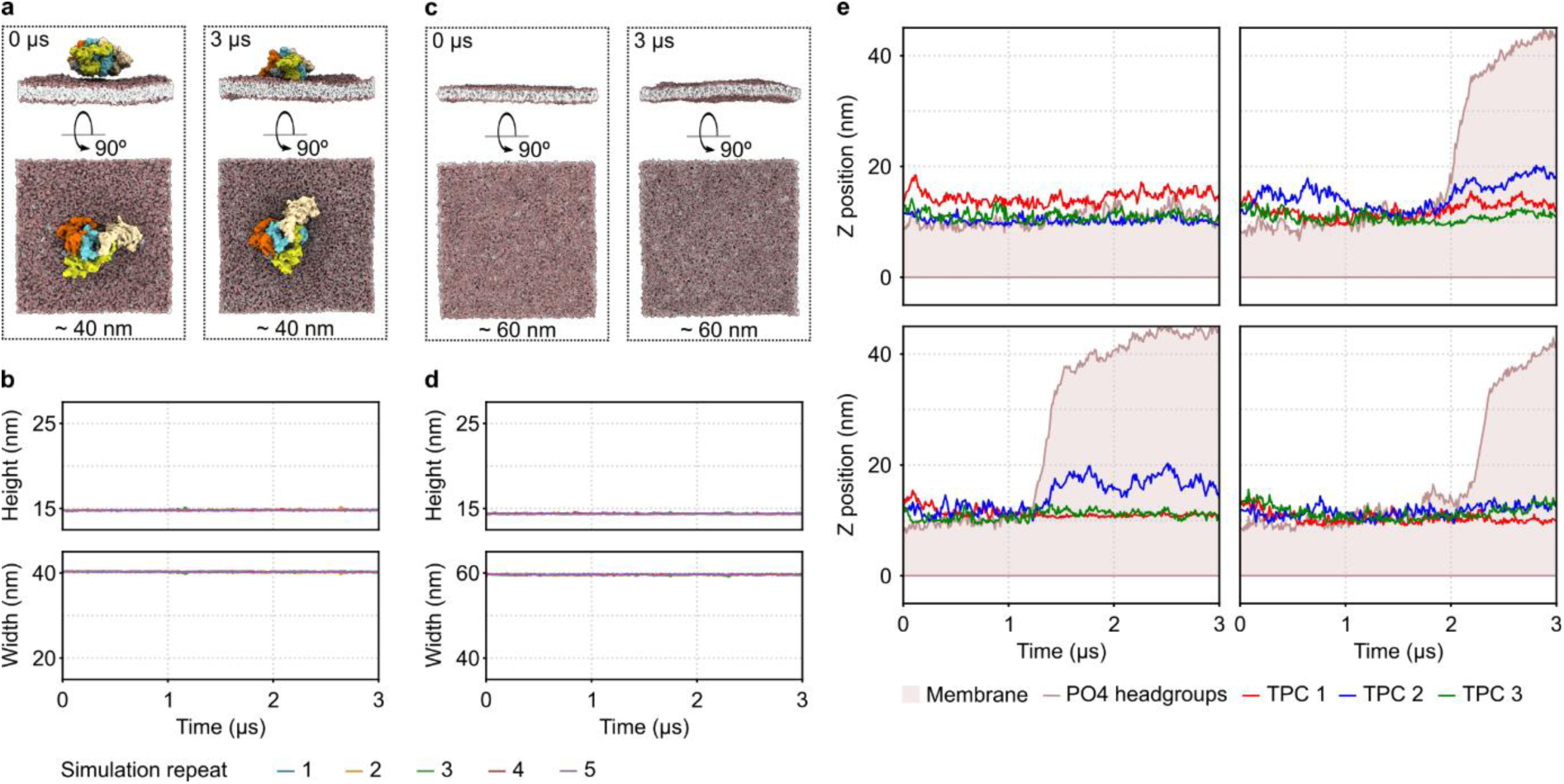
The structured domains of TPC provide the force for membrane deformation. a) Side and top view images of representative timepoints from the MD simulation of 40 nm x 40 nm membrane with a single TPC. The 0 µs time point represents the starting point and the membrane is roughly 40 nm x 40 nm in size. At 3 µs, the membrane has essentially the same size. b) Visualization of deformation of the simulation box over the simulation time for a 40 nm x 40 nm membrane with a single TPC. Membrane deformations do not occur. Different colors indicate different simulation repeats. c) Side and top view images of representative timepoints from the MD simulation of 60 nm x 60 nm membrane without TPC. The 0 µs time point represents the starting point and the membrane is roughly 60 nm x 60 nm in size. At 3 µs, the membrane has essentially the same size. d) Visualization of deformation of the simulation box over the simulation time for a 60 nm x 60 nm membrane without TPC. Membrane deformations do not occur. Different colors indicate different simulation repeats. e) Visualization of the membrane deformations along the Z axis (brown line and area) over the time alongside with the positions of the center of mass of the three TPC subunits (red, blue and green lines) for four MD simulation repeats not shown in Figure 5 d.

**Extended Data Fig. 10.**
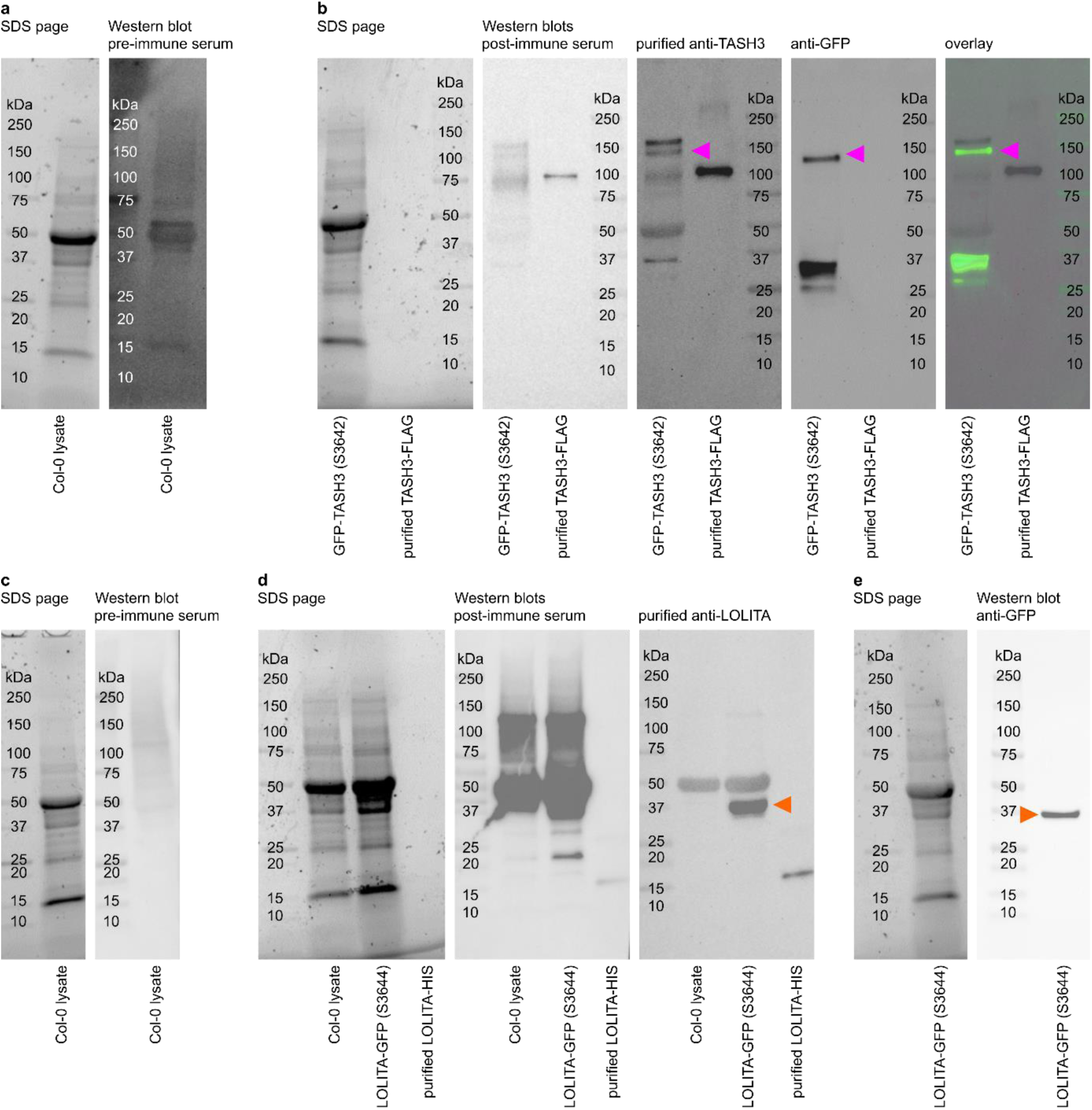
TASH3 and LOLITA antibody development. a) and c) The pre-immunization sera of the rabbits were tested against Col-0 lysate to ensure the absence of background immunity. b) The post-immune serum, as well as the purified anti-TASH3 antibody were tested against plant lysate of a GFP-TASH3 expressing plant, as well as against purified TASH3-FLAG. The serum generates a substantial amount of background signal. The purified anti-TASH3 antibody has less background signal in the plant sample, but it still recognizes multiple bands. It works well against purified samples. The specificity in the plant lysate was confirmed via anti-GFP (magenta arrow heads). d) The post-immune serum of the anti-LOLITA antibody showed a substantial amount of background signal in plant lysates, whereas the purified antibody presented only little background signal. The purified antibody was not sufficient to detect native LOLITA in Col-0 lysate, however it could detect both purified LOLITA-HIS and LOLITA-GFP expressed *in planta*. e) The specificity of the anti-LOLITA antibody was again confirmed against anti-GFP in the LOLITA-GFP expressing sample (orange arrow heads d & e).

**Extended Data Table 1 - Primers, plasmids, plant lines**

**Extended Data Table 2 - GFPtrap MS results**

**Extended Data Table 3 - XL-MS results**

**Extended Data Table 4 - Single band SDS page MS results**

**Extended Methods**

**Video 1: Integrative modeling of the TPC structure**

**Video 2: Structured domains of TPC deform the membrane**

### Extended Methods

#### Cloning/molecular work

Multiple methods were used to generate the plasmids used in this study. Genes were amplified via polymerase chain reaction (PCR) and then cloned via either Gibson assembly, Golden Gate cloning, or via Gateway BP and LR recombinations. To ensure compatibility with Golden Gate cloning, any BsaI restriction sites were mutated through silent mutations. All entry clones were sequenced. Expression clones were validated via restriction digest and by sequencing any recombination sites. An extensive list of primers and plasmids that were generated can be found in the Extended Data Table 1.

#### PSB-D cell culture work

To purify TPC from Arabidopsis PSB-D cell suspension cultures, Dark-grown cultures were transformed with recombinant TPLATE containing a C-terminal affinity tag consisting of a TEV cleavage site, GFP and a StrepTagII tag. The transformation and maintenance of the culture was done as described^1^. Cultures were selected via Kanamycin to guarantee successful transformation and the recombinant gene expression was monitored via anti-GFP Western Blot. Approximately every 6 months the cultures were re-transformed, because expression levels significantly reduced over time. One TPC purification required 10 L - 30 L (200 g - 600 g pellet) of PSB-D culture. To ensure enough material, the cultures were continuously upscaled and harvested by filtering the pellet. Afterwards the pellet was shock frozen and stored at −70 °C.

#### TPC Purification from PSB-D cultures

We designed a recombinant TPLATE allele with a TEV cleavage site followed by a GFP and TwinStrepTagII domain for TPC purification. This particular bait design utilizes the StrepTactinXT system from IBA life sciences^2^. In early StrepTag purifications followed by MS analysis we observed that the Arabidopsis proteome contains approximately 200 specific StrepTactin XT binding proteins (data not shown). Therefore, utilizing the TEV site to elute TPC specifically while contaminants remain bound to the beads, was key to a clean purified sample. Via the GFP domain we could track expression levels, as well as determine the cleavage efficiency during the purification by Western Blot. A weak naturally occurring IMAC binding site in the TWD40-2 subunit of TPC enabled us to then further clean up the complex and remove excess bait, as well as residing TEV (Extended Data Figure 1 a).

200 g - 600 g PSB-D culture pellets stored at −70 °C were ground to a very fine powder in liquid nitrogen first with a Braun MultiQuick 7 blender, then by hand with pestle and mortar. The powder was then dissolved and thawed in a lysis buffer (100 mM Tris/HCl pH 8, 150 mM NaCl, 0.5 mM TCEP, 1 tablet Roche Protease inhibitor/50 mL solution) with 1 mL of buffer per g of powder. Once thawed, Benzonase was added to a final concentration of 1 : 10 000 and the solution was incubated on ice for 30 minutes. Sonication could not be used for nucleic acid removal since the complex turned out to be sonication sensitive. After Benzonase digestion, the debris was removed via two subsequent centrifugations in 50 mL tubes at 45 000 x g at 4 °C for 20 minutes using a Sorvall Lynx II centrifuge. The lysate was transferred to clean tubes between the centrifugation steps. Following the centrifugation, the lysate was cleared via MiraCloth. The lysate was not filtered, because we observed high yield loss when filtering with different 0.45, or 0.22 micron filters. During the lysate centrifugation 1 mL of a 50 % suspension of StrepTactin XT 4flow HC beads was equilibrated, by washing it 4 times with lysis buffer. The equilibrated beads (0.5 mL bead volume) were added to the cleared lysate and then incubated for 3 h - 4 h at 4 °C while slowly rotating. After the incubation the beads were first harvested in 50 mL falcon tubes at 150g for 10 minutes at 4 °C, then they were collected in a 5 mL Eppendorf tube. To minimize losses, tubes were washed with a wash buffer (100 mM Tris/HCl pH8, 150 mM NaCl, 0.5 mM TCEP) two times and centrifuged again to capture any remaining beads. Once transferred to the 5 mL tube, the beads were centrifuged at 100 x g for 3 minutes at 4 °C. The beads were washed with the wash buffer at least six times with nine bead volumes (= 4.5 mL) per step. To elute TPC, one bead volume (= 0.5 mL) of wash buffer containing 0.1 mg/mL TEV was added and the beads were incubated rotating overnight at 4 °C. The next morning, the beads were washed twice with one bead volume (=0.5 mL) of the wash buffer and the supernatant of the two washing steps were combined with the elution fraction.

To remove excessive bait, as well as the TEV from our sample, we performed a subsequent IMAC. TPC has a weak naturally occurring IMAC binding site on its TWD40-2 subunit with the sequence HAHAHG, while the recombinant TEV provides a strong binding 7xHIS tag^3^. TPC elutes at 120 mM Imidazole present while the TEV remains bound to the IMAC beads at this concentration of Imidazole.

For the IMAC 100 μL - 50 μL of a 50 % Suspension of Ni Sepharose 6 Fast Flow beads (= 50 μL - 25 μL bead volume) was first rinsed with 10 bead volumes 500 mM Imidazole and then equilibrated in the wash buffer in a 1.5 mL Eppendorf tube, by washing 4 times with 20 bead volumes of the wash buffer per step. The equilibrated Ni Sepharose 6 Fast Flow beads were then added to the sample from the TEV elution and rotated for 2h at 4 °C. After incubation, the beads were harvested at 50g at 4 °C and washed three times with the wash buffer. Depending on the subsequent in vitro analysis the buffer was changed to accommodate for downstream experiments. For XL-MS experiments the beads were washed six times in a XL-MS reaction buffer (20 mM HEPES/NaOH pH 7.6, 250 mM NaCl, 50 mM Maltose, 0.5 mM TCEP) and for nsEM experiments the beads were washed in nsEM buffer 1 (50 mM HEPES/NaOH pH 7.6, 100 mM NaCl, 30 mM L-arginine, 0.5 mM TCEP), or nsEM buffer 2 (50 mM HEPES/NaOH pH 7.6, 100 mM NaCl, 30 mM creatine, 0.5 mM TCEP). TPC was then selectively eluted from the Ni-NTA Sepharose beads with 120 mM Imidazole in the corresponding buffer.

For XL-MS experiments, Imidazole was removed and the sample was concentrated to approximately 1 mg/mL by Vivaspin 500 50 kDa concentrator columns with a polyethersulfone (PES) membrane.

Protein concentrations were determined via Nanodrop and Qubit (ThermoFisher Scientific) and the sample quality was assessed via SDS page analysis. This purification protocol yields approximately 100 μg of TPC in total.

Since TPC proved to be unstable when stored at −70 °C under every condition tested, the protein was purified fresh for each downstream experiment.

#### SDS page analysis and Western blot

For protein gel electrophoresis samples were prepared by adding 4x Laemmmli Sample Buffer (product code: 1610747) with added NuPage reducing agent (Thermo Fisher NP0009). The samples were boiled for 8 minutes at 75 °C. The boiled samples were then loaded on a BioRad Mini-PROTEAN® 4-20 % stain-free gradient TGX gel. After running the samples at 180V for 40 minutes, the stain-free gels were developed using a BioRad ChemiDoc imaging system.

If Western Blot analysis was performed, the protein sample was transferred to a 0.2 μm PVDF membrane (BioRad) via the Trans-Blot turbo transfer system from BioRad. Afterwards the blots were developed with antibodies listed in the source file.

#### GFP trap MS experiments

The Miltenyi Biotec μMACS GFP isolation kit standard protocol for GFP trap experiments was used with adapted buffers to make sure TPC remained stable. Per pull down, 1 g of 5-day old Arabidopsis seedlings grown in constant light were ground in liquid nitrogen using pestle and mortar. The powder was dissolved on ice in an extraction buffer (20 mM HEPES/NaOH pH 7.6, 250 mM NaCl, 5 % Glycerol, 0.02 % DDM, 5 mM DTT, 1 tablet ROCHE protease inhibitor/50 mL) and once dissolved, the nucleic acids were digested by adding benzonase to a final concentration of 1 : 10 000 (v/v). After incubating for 30 minutes on ice, the debris was removed via centrifugation in 2 mL Eppendorf tubes at 18 000 x g for 30 minutes at 4 °C and the supernatant was cleaned with MiraCloth. The cleaned sample was then incubated for 2 h at 4 °C with 50 μL of anti-GFP μMACS beads and the standard protocol provided by Miltenyi Biotec was then followed. The wash buffer we used consisted of 20 mM HEPES/NaOH pH 7.6, 250 mM NaCl, 5 % Glycerol, 0.02 % DDM and 5 mM DTT. The final samples were eluted in 50 μL in 50 mM NH_4_HCO_3_ pH 8 for a downstream MS analysis.

The elution fractions from the GFP trap were reduced, alkylated and then trypsinated as described by^4^. The trypsinated peptides were then purified via Agilent C18 Omix tips (catalog number: A57003100) as described by the manufacturer.

Peptides were re-dissolved in 15 μL loading solvent A (0.1 % trifluoroacetic acid in water/acetonitrile (ACN) (98:2, v/v)) of which 5 μL was injected for LC-MS/MS analysis on an Ultimate 3000 RSLCnano system in-line connected to a Q Exactive mass spectrometer (Thermo). Trapping was performed on a 5 mM trapping column (Thermo scientific, 300 μm internal diameter (I.D.), 5 μm beads) in loading solvent A. After flushing from the trapping column, peptides were separated on a 25 cm Aurora Ultimate column (1.7 µm C18), with 75 µm inner diameter (Ionopticks) kept at a constant temperature of 45 °C. Peptides were eluted by a non-linear gradient starting at 1 % MS solvent B (0.1 % FA in water/acetonitrile (2:8, v/v)) reaching 33 % MS solvent B in 32 min, 55 % MS solvent B in 43 min, 70 % MS solvent B in 48 min, followed by a 2-minute wash at 70 % MS solvent B, and re-equilibration with MS solvent A (0.1 % FA in water) at a flow rate of 300 nl/min.

The mass spectrometer was operated in data-dependent mode, automatically switching between MS and MS/MS acquisition for the 5 most abundant ion peaks per MS spectrum. Full-scan MS spectra (400-2000 m/z) were acquired at a resolution of 70,000 in the Orbitrap analyzer after accumulation to a target value of 3,000,000. The 5 most intense ions above a threshold value of 13,000 were isolated with a width of 2.0 m/z for fragmentation at a normalized collision energy of 25 % after filling the trap at a target value of 50,000 for maximum 80 ms. MS/MS spectra (200-2000 m/z) were acquired at a resolution of 17,500 in the Orbitrap analyzer.

The Thermo raw files were processed using the MaxQuant software (version 2.2.0.0)^5^. Data was searched with the built-in Andromeda search engine against the Araport11plus database. This database contains all entries from the Araport11 database (The Arabidopsis Information Resource, www.arabidopsis.org/) concatenated with sequences of all types of possible contaminants in proteomics experiments. These include the **c**ommon **R**epository of **A**dventitious **P**roteins (cRAP) protein sequences, a list of proteins commonly found in proteomics experiments, present either by accident or by unavoidable contamination of protein samples (The Global Proteome Machine, www.thegpm.org/crap/). Additionally, commonly used tag sequences and typical affinity purification contaminants, such as sequences derived from the resins or the proteases used, were added. The Araport11plus database contains in total 49,058 sequence entries. To avoid misinterpretation and be able to distinguish between TML-linker and AP-2μ-linker, the TML-GFP and TML-AP-2μ-GFP sequences were split up in two common parts and two specific parts, as presented in Figure 4 and Extended Data Table 2, and added to the Araport11plus database.

Fixed modification was set to Carbamidomethylation of cysteines. Variable modifications were set to oxidation of methionines and N-acetylation of proteins N-termini. Mass tolerance on precursor ions was set to 4.5 ppm and on fragment ions to 20 ppm. Enzyme was set to Trypsin/P, allowing for 2 missed cleavages, and cleavage was allowed when arginine or lysine was followed by proline. PSM and protein identifications were filtered using a target-decoy approach at false discovery rate (FDR) of 1 %. Proteins identified with at least one unique peptide were retained.

The MaxQuant protein groups result file was uploaded in Perseus (version 1.6.15.0)^6^ for Label Free Quantification (LFQ) based differential analysis. Reverse hits, contaminants and only identified by site identifications were removed prior to uploading. LFQ intensity values were log2 transformed. Identifications were filtered for minimal three valid values in at least one group. Missing values were imputed with values around the detection limit, randomly drawn from a normal distribution with a width equal to 0.3 and a downshift equal to 1.8. Two-sided tests were performed, and significantly enriched proteins were determined by permutation-based FDR calculation, using thresholds FDR = 0.02 and S0 = 1. The results, including the imputed dataset, can be found in Extended Data Table 2.

#### XL-MS analysis

Cross-linking reactions were performed in solution with 13.5 μg TPC per reaction. For optimal cross-linking conditions we scouted different DSSO concentrations via SDS page and found 1.2 mM DSSO to be optimal (Extended Data Table 3).

First a 50 mM DSSO stock solution was prepared fresh by dissolving 1mg DSSO (Thermo Fisher A33545) in 51.5 μL DMSO. From this stock, a 10 mM solution was prepared on ice in the XL reaction buffer (20 mM HEPES/NaOH pH 7.6, 250 mM NaCl, 50 mM Maltose, 0.5 mM TCEP). Since DSSO is unstable in solution the cross-linking reactions were pipetted as fast as possible by mixing 5.1 μL reaction buffer, 2.4 μL DSSO 10 mM solution and 12.5 μL of purified TPC containing 13.5 μg protein. After incubating the 20 μL reaction at room temperature for 1h, the reaction was terminated by adding 180 μL of 50 mM NH_4_HCO_3_ pH 8.0.

To prepare the samples for MS, they were reduced with 5 mM DTT for 40 minutes at room temperature by adding 1 μL of 1M DTT stock solution. The reduced samples were alkylated 30 minutes at room temperature with 15 mM Iodoacetamide by adding 10 μL 0.3 M Iodoacetamide stock solution. Finally, the samples were digested with 1.25 μg Trypsin/LysC at 37 °C overnight by adding 5 μL of a 0.25 μg/μL Trypsin/LysC stock solution. To guarantee optimal digestion, the samples were digested for another 2 hours the next morning by adding 2.5 μL of a fresh 0.25 μg/μL Trypsin/LysC solution. The trypsinated peptides were dried via speedvac and then analyzed via LC-MS/MS.

Each sample was solubilized in 20 μL loading solvent A (0.1 % TFA in water:ACN (98:2, v:v)) moments before analysis. 5 μL was injected for LC-MS/MS analysis on an Ultimate 3000 RSLCnano system in-line connected to an Orbitrap Fusion Lumos mass spectrometer (Thermo). Trapping was performed at 20 μL/min for 1.5 min in loading solvent A on a µPAC^TM^ trapping column with C18-endcapped functionality (Thermo Scientific). The peptides were separated on a 50 cm µPAC™ column with C18-endcapped functionality (Thermo scientific). It was kept at a constant temperature of 50 °C. Peptides were eluted by a linear gradient reaching 4.5 % MS solvent B (0.1 % FA in water/acetonitrile (2:8, v/v)) after 9 min, 15 % MS solvent B at 120 min, 30 % MS solvent B at 150 min, 56 % MS solvent B at 173 min, followed by a 5-minutes wash at 56 % MS solvent B and re-equilibration with MS solvent A (0.1 % FA in water). The first 9 min the flow rate was set to 500 nl/min after which it was kept constant at 300 nl/min.

The mass spectrometer was operated in data-dependent mode. Full-scan MS spectra (375-1500 m/z) were acquired at a resolution of 60,000 in the Orbitrap analyzer after accumulation to a target AGC value of 2,800,00 with a maximum injection time of 60 ms. The precursor ions were filtered for charge states (3-8 required), dynamic exclusion (12 s; +/- 10 ppm window) and intensity (minimal intensity of 1.3E4). The most intense precursor ions during a cycle time of 1s were selected in the quadrupole with an isolation window of 1.5 Da and accumulated to an AGC target of 2E5 or a maximum injection time of 120 ms and activated using HCD fragmentation with stepped collision energy (NCE 27,30,33 %). The fragments were analyzed in the Orbitrap Analyzer at a resolution of 15,000.

Thermo raw files were analyzed using MS Annika 2.0^7^ integrated in Proteome Discoverer 2.4.1.15. For Merox 2.0^8^ analysis, raw files were first converted to mzML using MSConvertGUI v3.0.21084. Data analysis was performed against a database containing the 8 subunits of the TPLATE complex. The BS3 dataset from our previous publication^4^ was re-analyzed. The software specific settings and identified cross-links can be found in Extended Data Table 3. The cross-links were visualized using xiView^9^.

#### Single band SDS page MS analysis

The SDS page was stained with Colloidal Coomassie. Therefore, the page was incubated for 2h in fixing solution (50 % EtOH, 2 % H_3_PO_4_), washed 3x for 20 minutes with milliQ water and then incubated in a staining solution (34 % MeOH, 17 % (NH_4_)_2_SO_4_, 3 % H_3_PO_4_). While incubating in the staining solution, Coomassie Brilliant Blue G250 powder (Merck) was added to a concentration of 1 g/L. The page was incubated until the desired band saturation for excision was reached. The excised bands were transferred to 1.5 mL Eppendorf tubes and washed every 30 minutes with HPLC grade water until the gel plugs were completely destained. Then the samples were reduced by replacing the water with a reducing buffer (50 mM NH_4_HCO_3_ pH 8, 5 mM DTT) and shaking the samples for 40 minutes at room temperature. To alkylate, the reducing buffer was changed for an alkylation buffer (50 mM NH_4_HCO_3_ pH 8, 50 mM Iodoacetamide) and the samples were incubated for 30 minutes at room temperature. After alkylation the samples were washed two times with water before continuing with the in-gel trypsin digestion. First each sample was washed with 600 μL 95 % acetonitrile, then with 600 μL water and again with 600 μL 95 % acetonitrile. Each wash step was incubated for 10 minutes. Then the supernatant was removed and the shrunken gel plugs were placed on ice. 100 μL of digest buffer (50 mM NH_4_HCO_3_ pH 8, 10 % acetonitrile, 60 μg/mL Trypsin/LysC) was added and the samples were incubated on ice for 30 minutes and then digested at 37 °C for 3.5 hours. To release all digested peptides from the gel plugs, they were sonicated for 5 minutes. The supernatant with the trypsinated peptides was transferred to a clean tube and the plugs were washed with 50 μL replacement buffer (50 mM NH_4_HCO_3_ pH 8, 10 % acetonitrile). The wash fraction was added to the trypsinated peptides and the peptides were then purified via Agilent C18 Omix tips (catalog number: A57003100) as described by the manufacturer.

#### Polyclonal antibody generation

The antibodies against TASH3 and LOLITA were developed with Biotem. Synthesized peptides were used to immunize rabbits at day 0, 7, 14 and 34 after confirming that the pre-immune sera of the rabbits showed minimal response to Arabidopsis lysate samples via Western blot analysis. The peptide sequence used for LOLITA was KLGADNLKGVKNEEL+C and the peptide sequence for TASH3 was C+LEKVGDVPHKRKKGVFGTK. The peptides contained an extra cysteine (+C) to allow keyhole limpet hemocyanin conjugation as a carrier protein to the peptides. The rabbits were exsanguinated at day 42, after successful ELISA tests of the sera at day 28. The sera from day 42 were purified against the immunization target and the specificity of the polyclonal antibodies were tested against the target proteins. The SDS pages and Western blots confirming the antibody specificity and determining their background signal are depicted in Extended Data Figure 10.

#### Design of the TML-AP-2μ chimera

The TML-AP-2μ chimera construct was generated using Golden Gate cloning using fragments whose boundaries were defined via multiple sequence alignment.

The precise boundaries of the longin, the linker and the μ-homology domain of TML were determined via a multiple sequence alignment (MSA) using mafft v7^10,11^ following a blast^12–14^ search against all *Viridiplantae* with TML as an input sequence to generate a list of TML genes. Models of both the Arabidopsis TML sequence, as well as the MSA consensus TML sequence were calculated using the Robetta server via comparative modeling^15,16^. These models were then compared to AP-2 (PDBs 6QH5, 6QH6 and 6QH7)^17^. From this comparison all domains, as well as the disordered linker could be determined visually. The exact boundaries of the TML and AP-2 linkers were then chosen based on the sequence of the MSA and the chemical properties of the residues at the footprints of the linkers. After combining the TML longin domain, the AP-2μ linker and the μ-homology domain into one chimera, the design was modeled *in-silico* via Robetta to validate it. The integrity of both the longin, as well as the μ-homology domain were confirmed by comparing the Arabidopsis TML model with the chimeric model. The longin and μHD domain were PCR amplified and the linker sequence was ordered using primers (Extended Data Table 1). All fragments were combined via the GoldenGate system.

#### Plant line generation

The TMLp::TML-GFP, TMLp::TML-AP-2μ-GFP and H3.3p::TWD40-1-β-GFP plant lines were generated by floral dip^18^ of Col-0 plants or *tml-1* mutant plants^19^. Successful Arabidopsis transformants were either selected according to an antibiotic resistance, or according to a fluorescent reporter in the seeds, encoded by the introduced tDNA^20^. Positive selected transformants were then grown to the next generation, where they were selected for good heterologous gene expression and a segregation ratio of 3:1 for the transgene to ensure a single insertion. We selected at least two independent lines for each construct and confirmed the expression level of the introduced heterologous gene via Western blot.

For the colocalization analysis of the TML constructs with LAT52p::TPLATE-mSCARLET, the established homozygous TMLp::TML-GFP and TMLp::TML-AP-2μ-GFP expressing lines in the Col-0 background were transformed with a functional LAT52p::TPLATE-mSCARLET construct^21^. T1 transformants were selected via antibiotic resistance for LAT52p::TPLATE-mSCARLET and the T2 lines were imaged. Consequently, the imaged lines were homozygous for either TMLp::TML-GFP, or TMLp::TML-AP-2μ-GFP and heterozygous, or homozygous for LAT52p::TPLATE-mSCARLET.

To generate the plant lines used for the inducible depletion of PI4,5P_2_, dOCRL lines^22^ were transformed with the H3.3p::TWD40-1-β-GFP construct. T1 transformants were selected via fast red^20^ and imaged using a confocal microscope. Expression of the TWD40-1-β-GFP construct severely suffers from silencing in between generations, which prevented the establishment of stable homozygous lines. An overview of plant lines can be found in Extended Data Table 1.

#### Functionality test for TML constructs

The TMLp::TML-GFP and TMLp::TML-AP-2μ-GFP constructs were introduced in the *tml-1* mutant background and the transfer rate of the *tml-1* T-DNA insertion in the T2 generation was evaluated via the Sulfadiazine resistance located on the *tml-1* T-DNA^19^.

#### LUV flotation

Lipids (Avanti Polar Lipids) were dissolved in chloroform (PC, PE, and PA) or in chloroform, methanol, and water (PI4P and PI4,5P_2_). Lipids were mixed in ratios 80:20 for PC-PE, 60:20:20 for PC-PE-PA, 68:20:12 for PC-PE-PI4P, and 71:20:9 for PC-PE-PI4,5P_2_ and speedvac dried. The pellets were redissolved in 20 mM HEPES pH 7.5, 250 mM NaCl, 0.5 mM TCEP buffer and incubated for 90 minutes on a rotational wheel. Following the incubation, lipid suspensions were sonicated for 15 minutes in an ultrasonic bath. To obtain unilamellar liposomes of the homogenous size, the lipid suspensions were extruded through a polycarbonate membrane with a 0.2 μm pore size (Whatman).

For the co-flotation assays, 150 nmol liposomes per sample were mixed with purified TPC and incubated at room temperature for 45 minutes. After incubation, 100 μL of 60 % sucrose (w/v) was added to the sample resulting in a concentration of 30 % sucrose (w/v). The mixtures were transferred to centrifugation tubes, overlayed with 250 μL 25 % sucrose w/v and 50 μL 20 mM HEPES pH 7.5, 250 mM NaCl, 0.5 mM TCEP buffer and centrifuged for 2 hrs at 175,000 x g using S120-AT2 fixed angle rotor (Sorvall), 22 °C. After centrifugation, 250 μL bottom fraction (B), 150 μL middle fraction (M), and 100 μL top fraction (T) were collected with Hamilton’s syringe from the bottom of the tube. The pellet (P) was resuspended in 50 μL 20 mM HEPES pH 7.5, 250 mM NaCl, and 0.5 mM TCEP buffer.

Samples from individual fractions were mixed with 4× Laemmli loading buffer (Bio-Rad) and 4× XT Reducing Agent (Bio-Rad) and heated for 5 min at 70 °C. The samples were loaded on 4-20 % Mini-PROTEAN TGX Stain-Free Gels (Bio-Rad) and transferred to 0.2 μm PVDF membranes (Bio-Rad) using the Trans-Blot Turbo system (Bio-Rad). The membranes were blocked with 5 % nonfat dry milk (w/v) in TBS-T, followed by incubation with anti-TPLATE primary antibody and anti-rabbit secondary antibody (Sigma Aldrich). Clarity ECL Western Blotting Substrate (Bio-Rad) was used for visualization. The distribution of the TPC between the fractions was quantified based on band intensities in ImageJ software^23^.

#### LUV tubulation

Lipids (Avanti Polar Lipids) were dissolved in chloroform in the case of PC, PE, and PA and in chloroform, methanol, and water in the case of PI4P and PI4,5P_2_. 800 nmol lipids were mixed in ratios 80:20 for PC-PE, 60:20:20 for PC-PE-PA, and 71:20:9 for PC-PE-PI4,5P_2_ and subsequently speedvac dried. The pellets were redissolved in 500 μL 20 mM HEPES, 15 % (w/v) raffinose, 0.5 mM TCEP and incubated for 90 minutes on the rotational wheel. Following the incubation, lipid suspensions were sonicated for 15 minutes in an ultrasonic bath. To obtain unilamellar liposomes of homogenous size, lipid suspensions were extruded through a polycarbonate membrane with a 0.2 μm pore size (Whatman). Liposomes were diluted up to 1 mL with 20 mM HEPES, 250 mM NaCl, 0.5 mM TCEP buffer, pH 7.5 and centrifuged 30 min at 74000g using S120-AT2 fixed angle rotor (Sorvall), 22 °C. The supernatant was discarded, the pellet was resuspended in 500 μL 20 mM HEPES, 250 mM NaCl, 0.5 mM TCEP buffer, pH 7.5 and the centrifugation step was repeated. The supernatant was discarded, and the pellet was resuspended in 100 μL 20 mM HEPES, 250 mM NaCl, 0.5 mM TCEP buffer.

For the tubulation reaction the liposomes were diluted to a final concentration of approximately 2 mM. To find the optimal protein concentrations, several dilutions of TPC were tested. 10 μL of freshly purified TPC in tubulation buffer (20 mM HEPES/NaOH pH 7.6, 250 mM NaCl, 50 mM Maltose, 0.5 mM TCEP) were gently mixed with 20 μL of liposomes and incubated for 45 minutes on a rotating wheel at 6 rpm. For the no protein control, 10 μL of tubulation buffer was added to 20 μL of liposomes. Afterwards the samples were diluted 4x in the tubulation buffer and used right away for negative stain EM grid preparation.

While the tubulation reaction was incubating, Formvar FCF400-CU-50 EM grids were activated via glow discharging with 5mA for 20s in an ELMO glow discharger. The glow discharged grids had to be used within 60 minutes. 10 μL of the sample were applied per grid for 1 minute and otherwise the standard negative staining EM procedure with uranyl acetate was followed.

The micrographs were then imaged on a JEOL 1400+ microscope with a LaB6 filament at 120kV acceleration voltage using a TVIPS F416 CCD camera at 25 000x magnification. The CS value equals 3.4 and the calibrated pixel size corresponds to 4.78Å/px at 25 000x magnification.

#### Confocal microscopy

Colocalization analysis between TPLATE and TML was performed on a Nikon Ti microscope with the Perfect Focus System (PFSIII) for Z-drift compensation, equipped with an Ultraview spinning-disk system (PerkinElmer) and two 512×512 Hamamatsu ImagEM C9100-13 EMccd cameras. The microscope was controlled via the Volocity V7.0.0 software package. IDePP imaging was performed on the spinning disk and on a Leica DMI8 TCS SP8X confocal microscope with a white light laser and hybrid detectors (HyD). Imaging of TWD40-1-β-GFP in combination with FM4-64 was performed on the Leica SP8X.

5 day old seedlings of the iDePP control line expressing UBQ10pro:mCIT-2xPHPLC/pB7 (mCITRINE-2xPH) as a PI4,5P_2_ marker and the dexamethasone (DEX) inducible PI4,5P_2_ phosphatase BQ10pro:GVG-tE9::6xUAS-35Smini:MAP-mCH-dOCRL168-509/pH7 (MAP-mCherry-dOCRL)^22^ were imaged after 0h and 16h 5 μM DEX induction. The seedlings were imaged with a 60x water immersion objective on the Perkinelmer spinning disk system. mCITRINE was excited with a 514 nm diode laser at 40 % power and the signal was recorded between 525 nm to 575 nm. mCherry was excited using a 561 nm diode laser at 60 % power and the signal was recorded between 580 nm to 650 nm. Both channels were imaged sequentially using single camera mode with 500 ms exposure time.

For the colocalization analysis between TPLATE-mSCARLET and either TML-GFP, or TML-AP-2μ-GFP, the plant lines were imaged and the data was analyzed as described in our previous publication^24^. To control for random associations of fluorescent signals at the PM, the GFP channel was rotated by 90° to the right and then re-analyzed. Therefore, 3–5-day old, etiolated hypocotyl epidermal cells were imaged with an 100x oil immersion objective on the PerkinElmer spinning disk system. GFP was excited with a 488 nm laser and the emission was recorded between 500 nm to 550 nm, while mSCARLET was excited with a 516 nm laser and the emission was recorded between 545 nm to 685 nm. The exposure time was 750 ms and 2-minute-long videos were recorded in dual camera mode after aligning both cameras.

The TWD40-1-β-GFP lines stained with FM4-64 and TWD40-1-β-GFP lines inducibly co-expressing MAP-mCherry-dOCRL^22^ were imaged on a Leica DMI8 TCS SP8x confocal microscope with a white light laser and a hybrid detector (HyD) with gating functionality. 5-day old seedlings were imaged with a 63x water immersion lens. The green and red signals were recorded sequentially. TWD40-1-β-GFP was excited using a white light laser with three-line excitation at 472 nm, 480 nm and 488 nm simultaneously with 70 % laser power and the signal was detected between 500 nm - 525 nm with a HyD detector with gating. For FM4-64-stained seedlings, the dye was excited at 515 nm and the emission was detected between 590 nm - 725 nm with a HyD detector. Lines expressing MAP-mCherry-dOCRL were excited at 582 nm and the emission was detected between 595 nm - 675 nm with a HyD detector with gating.

#### Negative staining of EM grids

Formvar FCF400-CU-50 EM grids were glow discharged for 30 seconds with a 5mA current using an ELMO glow discharger. The grids were handled using Dumont N5 negative action style tweezers. For the staining three 20 μL drops of 2 % uranyl acetate solution were prepared on a parafilm. 4 μL of purified TPC was applied per glow discharged grid. After 30 seconds the protein was blotted away using Whatman paper No. 1 and the grid was immediately placed on top of the first uranyl acetate drop for 10 seconds. Afterwards the uranyl acetate was removed via Whatman paper No. 1 and the grid was placed on the second uranyl acetate drop for 1 second. After blotting away the uranyl acetate the grid was incubated for 1 minute on the third uranyl acetate drop. Finally, the uranyl acetate was blotted away, leaving a thin film and the grids were dried at room temperature before imaging.

#### TPC negative stain EM dataset acquisition

Negative stain grids with purified TPC were imaged on a JEOL 1400+ microscope with a LaB6 filament at 120 kV acceleration voltage using a TVIPS F416 CCD camera at 60 000x magnification. The CS value equals 3.4 and the calibrated pixel size corresponds to 1.94Å/px at 60 000x magnification. Automated data acquisition was performed via Serial EM v4.0.28^25,26^. For high throughput multi shot imaging via beam shift the usual calibrations and corrections were performed, like astigmatism correction and coma free alignment. The dataset was then acquired in a defocus range of −0.5 μm --3 μm with a step size of 0.5 μm. Before data processing, micrographs with visible artifacts, like drift and protein aggregation were discarded. The following 3D map generation was done with CryoSparc.

#### 3D map generation of negative stain dataset via Cryosparc

The nsEM datasets were evaluated with CryoSparc v4.1.2^27^. The micrographs recorded by SerialEM were converted from 16bit tiff to mrc file format via IMOD v4.11^28,29^. Afterwards the images were imported to our CryoSparc server and CTF estimation was performed via CTFFIND4 v4.1.14^30^. After CTF estimation bad micrographs were removed from the dataset. Bad micrographs were micrographs that showed astigmatism, or drift (visible in the thone rings), or where CTF estimation failed (low contrast, artifacts, etc). About 1.7 % of micrographs were removed in this step resulting in 1 698 good micrographs for our full L-arginine dataset. From these micrographs particles were automatically picked using the pre-trained topaz resnet8 model with 64 units, a downsampling factor of 8 and a radius of 8 with topaz v0.2.5^31^. Since negative stain EM micrographs generally present white noise as a consequence from the staining, the set of picked particles from topaz contained artifacts. Therefore, the final set of particles was selected in three rounds of 2D classification using a maximum resolution of 15 Å, a circular mask diameter of 270 Å (resulting in a box size of 324, equalling roughly 1.5x the longest axis of TPC) and a white noise model. During the first two rounds artifacts and clearly bad classes were discarded, while in the last round only good 2D classes were selected. This 2D classification yielded a particle set of 156 997 good particles for the full L-arginine dataset. To ensure no good particles were removed during 2D classification, all discarded particles were re-classified (Extended Data Figure 1 e). The final set of particles was then used for ab-initio 3D map generation where we limited the maximum resolution to 15 Å, otherwise default CryoSparc parameters were used. Since 3D classification only yielded one 3D class for our dataset, we refined the ab-initio map using all particles. For 3D refinement CryoSparc’s homogenous refinement was used with a maximum align resolution of 11 Å, otherwise CryoSparc over-fitted the refined map significantly. The remaining parameters were left as default. The reconstructed map for the full L-arginine dataset achieved a resolution of roughly 17 Å (Extended Data Figure 1 g) and presented a volume corresponding to a protein complex of 1MDa. It also showed preferred orientations, as is common with negative stain EM (Extended Data Figure 1 d & i). Therefore, to avoid artifacts we gaussed the map to approximately 20 Å resolution before using them for our integrative approach.

#### Experimental validation of the generated 3D map

To validate the obtained negative stain EM map of TPC, we split the L-arginine dataset recorded from protein purified in 50 mM HEPES/NaOH pH 7.6, 100 mM NaCl, 30 mM L-arginine, 0.5 mM TCEP in two parts based on the grid they were recorded from. Part 1 consisted of 681 micrographs and part 2 consisted of 1 017 micrographs. Then we reconstituted 3D maps as described above independently for each part. For part 1 we yielded a particle set of 64 520 good particles, for part 2 the particle set contained 91 720 particles. In addition to that we also recorded a creatine dataset of TPC purified in 50 mM HEPES/NaOH pH 7.6, 100 mM NaCl, 30 mM creatine, 0.5 mM TCEP and reconstituted a 3D map of this dataset. Here the final particle set consisted of 73 355 particles extracted from 1 190 micrographs. All reconstructed maps had the same shape, achieved a resolution of roughly 17 Å and presented a volume corresponding to a protein complex of 1MDa. They also showed the same preferred orientations. The maps were compared using UCSF ChimeraX^32,33^. Therefore, the volumes were adjusted to match exactly 1MDa. Afterwards pairwise comparisons were calculated using the fit in map command (Extended Data Figure 1 c).

#### ConSurf analysis

To determine structural conservation, we analyzed the protein sequences of interest with a local installation of ConSurf^34^. The results can be found in the source file. To generate Extended Data Figure 5 we simplified the ConSurf output by calculating a floating average of ConSurf scores considering 5 residues. Afterwards we assigned the color green to a ConSurf score below 5 resembling a variable sequence and purple to a score above 5 representing conserved regions.

#### Integrative structure modeling of the TPC

Python modeling interface of the integrative modeling platform (IMP) package version 2.17^35,36^ was used to build the TPC structure. The representation of the individual TPC subunits was derived from a structure of the given subunit deposited in the Alphafold2 database^37^. Well-structured (reliably predicted) domains of the TPC subunits were represented by beads of varying sizes, 1 to 10 residues per bead, arranged into a rigid body. Loop regions were represented by a flexible string of beads of corresponding size to the respective rigid body. Regions predicted with very low confidence were represented by a flexible string of beads corresponding to 20 residues each. The TML linker (region from 186 to 406) was represented by beads corresponding to 10 residues each. Overall, the system consisted of 12 rigid bodies and 229 flexible beads. Alphafold-multimer, as implemented in the ColabFold notebook^38^, was used to model the TPC tetrameric core composed of LOLITA, the TASH3 trunk domain, the TPLATE trunk domain and the TML longin domain, and the dimer of TWD40-1 and TWD40-2. The initial structure of the TPC hexamer was obtained by superimposing the TPC tetrameric core and the TWD40-1/TWD40-2 dimer on the structure of COPI (PDB code 5a1u). The assembled TPC hexamer was fitted into the nsEM map represented as a Gaussian mixture model (GMM)^39^ using UCSF ChimeraX^32,33^. Other structural parts of TPC were randomly positioned around the GMM representation of the nsEM map. 123 and 126 unique intra- and intermolecular BS^3^ and DSSO cross-links were used to construct the scoring function. In addition to the cross-link datasets, the excluded volume restraints, the sequence connectivity restraints and the GMM map were added to the scoring function. Replica Exchange Gibbs Monte Carlo algorithm^40^ was used to sample structures satisfying input restraints. The sampling produced a total of 720,000 models from 24 independent runs. Models were clustered based on cross-linking and GMM agreement, sequence connectivity, excluded volume, and total score. The four-step protocol tested the top-scoring cluster for convergence, exhaustiveness, and precision^41^. The PrISM method^42^ was used to analyze the precision of individual regions of the integrative TPC model. Alphafold3 server^43^ was used to produce a model of the truncated TPC hexamer.

#### Molecular dynamics simulation

All simulations were performed using the GROMACS software^44^. The integrative structural model of TPC was used as an input for MD simulations. Missing loops in the rigid body representation of the TPC subunits were modeled using MODELLER software^45^. To relax the TPC structure, a short atomistic MD simulation was performed using CHARMM36m force-field^46^. Using the CHARMM-GUI web server^47,48^, TPC was placed in a 25×25×25 nm simulation box solvated with 0.15 M KCl in water and neutralized. The system was energy minimized using the steepest descent algorithm up to the maximum of 5000 steps and equilibrated for 125 ps. During energy minimization and equilibration, position restraints were introduced on protein backbone and side chains. 10 ns production run was performed with a 2 fs time step in the NPT ensemble. Pressure was set to 1 bar and maintained with the Parrinello-Rahman barostat, with a coupling constant of 5.0 ps and compressibility of 4.5e-4. Temperature was set to 303.15 K and maintained with the Nose-Hoover thermostat with a coupling constant of 1.0 ps.

MARTINI 2.2 coarse-grained (CG) force-field^49^ was used for the simulations of TPC with a membrane. The relaxed TPC structure was converted to coarse-grained (CG) representation using the CHARMM-GUI web server^47,48^. AtEH1, AtEH2, TASH3_1-101_, the SH3 domain and the TWD40-1 all-alpha domain were removed for the purpose of CG MD simulations. Elastic Network in Dynamics (ElNeDyn) was applied to retain the secondary and tertiary structure ^50^.

For the simulations of a single TPC, a 25×25 nm membrane composed of 50 % PC, 50 % PE in the upper leaflet and 36 % PC 36 % PE 10 % PS 10 % PA 5 % PI4P and 3 % PI4,5P_2_ in the lower leaflet was setup using insane.py script^51^. The charge of PA was modified to −2. TPC was positioned in the proximity of the pre-equilibrated membrane patch and inserted in a 25×25×25 nm simulation box. Subsequently, the systems were solvated with 0.15 M NaCl in water and neutralized. The system was energy minimized using the steepest descent algorithm up to the maximum of 5000 steps and equilibrated for 4750 ps. Position restraints on protein backbone and side chains were progressively released during equilibration. 5 production runs, 5 μs each starting from different initial velocities were performed with a 20 fs time step in the NPT ensemble. Pressure was set to 1 bar and maintained with the Parrinello-Rahman barostat, with a coupling constant of 12.0 ps and compressibility 3.4e-4. Temperature was set to 323 K and maintained with the v-rescale thermostat with a coupling constant of 1.0 ps. The bond lengths were constrained using the LINCS algorithm. Van der Waals and Coulomb cut-offs were set to 1.1 nm.

Per residue-contacts with lipids were analyzed using the PyLipID software^52^. Last 2 μs from the 4 trajectories, in which TPC interacted with the membrane through the membrane, were selected for contacts calculations. Such trajectories were identified by calculating the minimum distance between TWD40-1, TWD40-2, and TASH subunits and the membrane using a GROMACS in-build tool, *gmx mindist.* PO4 and CNO headgroup beads were used for PS, PO4 for PA, PO4 and P1 for PI4P and PO4, P1 and P2 headgroup beads for PI4,5P_2_ with the distance cut-off 0.7 nm. The contacts were visualized as occupancies mapped on TPC structure using UCSF ChimeraX software^32,33^. Trajectories were further analyzed using GROMACS in-build tools, *gmx rotmat* and *gmx densmap,* respectively. Membrane curvature was analyzed using the MembraneCurvature toolkit implemented in the MDAnalysis software^53,54^. Gaussian interpolation was used to visualize the results.

For the simulations with asymmetric membranes, a 40×40 nm membrane was set up using insane.py script^51^. The membrane was composed of 50 % PC, 50 % PE in the upper leaflet and 36 % PC 36 % PE 10 % PS 10 % PA 5 % PI4P and 3 % PI4,5P_2_ in the lower leaflet. The lipid count was asymmetric, with the lower leaflet containing 1.2 times more lipids than the upper leaflet. The charge of PA was modified to −2. 1x TPC was positioned in the proximity of the pre-equilibrated membrane patch and inserted in a 40×40×25 nm simulation box. Conditions of solvation, energy minimization and production run were as previously described. 5 production runs of 3 μs each starting from different initial velocities were performed.

Further, a 60×60 nm membrane was set up using insane.py script^51^. The membrane was composed of 50 % PC, 50 % PE in the upper leaflet and 36 % PC 36 % PE 10 % PS 10 % PA 5 % PI4P and 3 % PI4,5P_2_ in the lower leaflet. The lipid count was asymmetric, with the lower leaflet containing 1.2 times more lipids than the upper leaflet. The charge of PA was modified to −2. 3x TPC were positioned in the proximity of the pre-equilibrated membrane patch and inserted in a 60×60×25 simulation box. Conditions of solvation, energy minimization and production run were as previously described. 5 production runs of 3 μs each starting from different initial velocities were performed.

Finally, for the simulations of the membrane without TPC, the above described 60×60 nm membrane was inserted inserted in a 60×60×15 nm simulation box. Conditions of solvation, energy minimization and production run were as previously described. 5 production runs of 3 μs each starting from different initial velocities were performed.

The width and height of the simulation box in time were calculated using a GROMACS in-build tool *gmx energy.* Z positions were analyzed using the MDAnalysis software^53,54^. By tracing the highest and lowest z-positions of the phosphate (PO4) headgroup beads, we monitored the membrane budding events. By tracing the z-positions of the center of mass of each of the three TPCs, we monitored the position of the protein towards the bud. To plot the z-positions of the individual components, all the values were translated so that the lowest point of the upper leaflet z=0. Height and width of the simulation box in time were analyzed using a GROMACS in-build tool *gmx energy*.

#### Visualization of protein structures and data

For the visualization of all protein structures, UCSF ChimeraX^32,33^ was used. All figures were prepared with the Inkscape program (https://inkscape.org/).

#### Statistical analysis

The R package in R studio was used for statistical analysis (https://www.R-project.org/). For multiple comparison the multcomp package was used^55^.

## Notes

### Competing Interest Statement

The authors have declared no competing interest.

